# Estimating Assembly Base Errors Using K-mer Abundance Difference (KAD) Between Short Reads and Genome Assembled Sequences

**DOI:** 10.1101/2020.03.17.994566

**Authors:** Cheng He, Guifang Lin, Hairong Wei, Haibao Tang, Frank F White, Barbara Valent, Sanzhen Liu

**Author notes:** Corresponding authors: Sanzhen Liu, Department of Plant Pathology, Kansas State University, 4024, Throckmorton Center, Manhattan, Kansas, 66506.

## Abstract

Genome sequences provide genomic maps with a single-base resolution for exploring genetic contents. Sequencing technologies, particularly long reads, have revolutionized genome assemblies for producing highly continuous genome sequences. However, current long-read sequencing technologies generate inaccurate reads that contain many errors. Some errors are retained in assembled sequences, which are typically not completely corrected by using either long reads or more accurate short reads. The issue commonly exists but few tools are dedicated for computing error rates or determining error locations. In this study, we developed a novel approach, referred to as K-mer Abundance Difference (KAD), to compare the inferred copy number of each k-mer indicated by short reads and the observed copy number in the assembly. Simple KAD metrics enable to classify k-mers into categories that reflect the quality of the assembly. Specifically, the KAD method can be used to identify base errors and estimate the overall error rate. In addition, sequence insertion and deletion as well as sequence redundancy can also be detected. Therefore, KAD is valuable for quality evaluation of genome assemblies and, potentially, provides a diagnostic tool to aid in precise error correction. KAD software has been developed to facilitate public uses.

## INTRODUCTION

DNA sequencing technologies have revolutionized genetic and genomic analyses, facilitating *de novo* assemblies of genomes from various species with small to large complex genomes of species such as wheat (1). Genome assemblies using Illumina technologies, here referred to as short-read sequences, are typically highly fragmented but sequence bases are accurate. The contiguity of genome assemblies can be dramatically improved by using long single-molecule sequencing technologies led by two technologies, principally by Pacific Biosciences SMRT and Oxford Nanopore (ONT) platforms (2). Sequencing reads yielded from both technologies have relatively high rates of errors, and are dominated by small insertions or deletions. Although consensus sequences from high coverage of sequencing reads reduce errors in genome assemblies, nonrandom errors, or biased errors, in reads can result in inaccurate consensus sequences. Biased errors could be caused by epigenomic modifications that affect sequencing signals for base calling (3, 4).

To mitigate per base errors in a draft assembly, sequencing polishing algorithms have been developed to mitigate such issues through using signal-level raw data (5, 6). However, for genome sequences of many species, a great number of errors still exist after multiple rounds of sequence polishing, or error correction (7). Practically, errors can be further reduced via correction with additional Illumina short reads. Variants revealed by alignments of Illumina reads with assembled sequences are typically used for error correction (8). This strategy works well for small low-repetitive genomes because most assembly regions can be uniquely covered by Illumina reads. For large repetitive genomes, the strategy works less well due to a lower proportion of genomes uniquely aligned by Illumina reads, a higher rate of misalignments, and even no alignments at some poorly assembled regions (9).

Assembly quality of genome sequences is related to assembly contiguity and completeness, correctness of sequence ordering, and consensus base accuracy. Community-based projects like GAGE (10) and Assemblathon (11) looked at a suite of criteria for a comprehensive assessment when benchmarking different assembly methods when the ground truth assembly is known. The value of N50, the length of the smallest contig of a set of the top long contigs that cover half of assembly space, is widely used as an indicator of assembly contiguity. Alignment rates or number of variants based on alignment to assembled sequences with genome sequencing reads, RNA sequencing reads, or comparison against the reference genomes of related varieties or species can indicate assembly quality. Tools were developed for comparing some of these parameters for genome assemblies, such as Quast (12). Conserved benchmarking universal single-copy orthologs (BUSCO) (13) or core eukaryotic genes mapping approach (CEGMA) (14) were used to assess genome completeness simply based on evaluating the coding or gene space. LTR Assembly Index (LAI) that indicates the assembly quality of LTR (Long Terminal Repeats) retrotransposons was designed to evaluate assembly continuity, extending assembly quality assessment to repetitive regions (15).

In addition, approaches for genome assembly characterization were also developed based on profiles of k-mers, substrings of length *k* from longer DNA sequences. K-mer based approaches have been used to quantify genome size, repetitive levels, heterozygosity in assembled sequences (16–19), and to perform reference-free genome comparisons based on sequencing data (20, 21). A method KAT (k-mer analysis toolkit) was developed to profile k-mer spectra of both sequencing reads and assemblies and to visualize difference of k-mer abundance in the assembly and read data (22). Here, we quantified abundance of k-mers from sequencing reads and k-mer occurrence in the assembly genome, and developed a single value, K-mer Abundance Difference (KAD), per k-mer. Given a set of input reads, KAD analysis can evaluate the accuracy of nucleotide base quality at both genome-wide and single-locus levels, which, indeed, is appropriate, efficient, and powerful for assessing genome sequences assembled with inaccurate long reads.

## METHODS

### Simulation of genome sequences with single nucleotide substitution errors

Genome sequences were simulated with various single nucleotide substitution errors using the software simuG (23). The parameters of simuG were set as “-snp_count <total nucleotide number of genome x variation rate> -titv_ratio 0.5”.

### Simulation of genome sequences with different error types

The software simuG was also used to simulate genome sequences with different error types (23). For single nucleotide substitution errors, the parameters of simuG were set as “-snp_count <total nucleotide number of genome * variation rate> -titv_ratio 0.5”. For short INDELs, the parameter was set as “-indel_count <total nucleotide number of genome × variation rate> -ins_del_ratio 1”. For long sequence redundancy, the parameter was set as “-cnv_count <number of long sequence redundancy> -cnv_gain_loss_ratio Inf”. For long sequence deletion, the parameter was set as “-cnv_count <number of long sequence redundancy> -cnv_gain_loss_ratio 0”.

### Simulation of reads with and without errors

The software DWGSIM (https://github.com/nh13/DWGSIM) was used to generate reads with or without errors (error-free reads) using reference genomes of multiple species with various genome sizes. To simulate reads without errors with different read depths, the parameter was set as “-e 0 -E 0 -C <read depth> −1 150 −2 150 -r 0 -R 0 -X 0 -y 0 -c 0 -S 0”. To simulate reads with single nucleotide substitution errors with 50x read depth, the parameter was set as “-e <sequencing error rate> -E <sequencing error rate> -C 50 −1 150 −2 150 -r 0 -R 0 -X 0 -y 0 -c 0 -S 0”. The parameter -C represents the read depth, the parameters −1 and −2 represent the lengths of the first and the second reads of paired-end reads. To simulate reads without errors, all the parameters control the error rates in reads (-r, -R, -X, -y, -c, -S) were set to 0.

### Calculation of True Capture Rate (TCR) and False Capture Rate (FCR)

Error k-mers were identified with the KAD script “KADprofile.pl”, which determined the KAD value per k-mer and identified error k-mers. To calculate the TCR value that stands for the percentage of simulated base errors detected by KAD, all the error k-mers detected by KAD were aligned to their simulated genomes with bowtie (24) and the overlapping error k-mers were merged into error regions, which was implemented by a KAD script “KADdist.pl”. The ratio of simulated errors located in error regions to total simulated errors was calculated as the TCR value. The FCR value stands for the percentage of error k-mers that do not overlap with simulated base errors. Therefore, the FCR value was determined by calculating the ratio of the number of error k-mers overlapping with simulated errors to the total number of error k-mers detected by KAD.

### Xvv1601 whole genome sequencing via PacBio and genome assembly

Bacterial growth and DNA extraction referred to a previous procedure (25). The 10-20kb whole genome shotgun libraries were constructed using Xvv1601 genomic DNAs. The library was sequenced with P6-C4 chemistry on a SMRTcells of PacBio RS II at the Yale Center for Genomic Analysis (YCGA). PacBio genome assembly: Canu (v1.3) was used for genome assembly (26). PacBio reads with the minimum length of 5 kb were used.

To generate Illumina data for the bacterium XV1601, the sequencing library was prepared using the Illumina TruSeq DNA LT sample Prep kit. Paired-end 2×300 bp reads were generated on an Illumina MiSeq at the Integrated Genomic Facility at Kansas State University. To examine the impact of read depths on error detection, the module “sample” of the software seqtk (https://github.com/lh3/seqtk) was used to down-sample Illumina reads to approximately 90x, 80x, 70x, 60x, 50x, 40x, 30x, and 20x.

### Identification of polymorphisms between two Xvv1601 assembly versions

The software MUMmer 4 (27) was used to identify DNA polymorphisms between the two assemblies (canu and final) of the bacterial strain Xvv1601. The two assembly sequences were aligned with the nucmer command. Alignments were filtered with the command delta-filter with (-1 -l 10000 -i 90) which resulted in unique alignments with at least 10 kb matches and at least 90% identity between the two assembled genomes. The alignments passing the filtering were used for the variant discovery with “show-snps”.

### B71 whole genome sequencing using Nanopore MinION

B71 nuclear genomic DNA was prepared as described previously (28). Genomic DNA was subjected to 20kb size selection using Bluepippin cassette kit BLF7510 with High-Pass protocol (Sage Science, USA), followed by library preparation with the SQK-LSK109 kit (Oxford Nanopore, UK). Library was loaded to the flowcell FLO-MIN106D (Oxford Nanopore, UK) and sequenced on MinION (Oxford Nanopore, UK). Guppy version 2.2.2 was used to convert Nanopore raw data (fast5) to fastq data with default parameters.

### B71 Nanopore genome assembly and sequence polishing

Nanopore reads were input to Canu 1.8 for genome assembly with the following parameters (genomeSize=45m minReadLength=5000 minOverlapLength=1000 corOutCoverage=80) (26). Nanopore reads were aligned back to the Canu assembly with minimap2 (2.14-r892) with the parameter of (-ax map-ont). Alignments in BAM format converted by Samtools (1.9) were input for assembly polishing with Nanopolish (version 0.11.0) with default parameters (https://github.com/jts/nanopolish). The Nanopolish procedure was repeated twice. Nanopolish-polished sequences were further polished with trimmed reads of Illumina sequencing data (SRA accession: SRR6232156) using Pilon (version 1.23) (8). In the Pilon polishing, Illumina reads were aligned to Nanopolish-polished sequences with the aligner bwa (0.7.12-r1039) with default parameters of the module “mem” (29). Pilon used bwa alignments and polish assembled sequences with the parameters of (--minmq 40 --minqual 15). Assembled contigs were renamed based on their similarity to the assembly of B71Ref1 (28). We also manually fixed a misassembly that joined a previously identified mini-chromosome’s sequence with chromosome 6. The assembly and polishing procedure resulted in ONTv0.14. The same Pilon procedure was applied to further polish the PacBio assembly B71Ref1 (28) with Illumina reads, resulting in a new assembly PBRef1.3.

### Whole genome sequence alignment via NUCmer

The nucmer command from the software MUMmer 4 (27) was used for whole genome alignment between B71Ref1.3 and ONTv0.14. The parameter of “-L 1000” was used in the command nucmer and the parameter of “-L 5000 -I 98” in the command of show-coords, which resulted in alignments with at least 5 kb matches and at least 98% identity between the two assemblied genomes.

### KAD analysis to analyze B71 genome assemlies

Using trimmed B71 Illumina reads, the KAD analysis was performed for both assemblies ONTv0.14 and B71Ref1.3 with the script “KADprofile.pl”, which determined the KAD value per k-mer and grouped k-mers. The script “KADdist.pl” was used to map k-mers to the assembly genomes and profile distributions of k-mers from each k-mer group, particularly the group of error k-mers.

## RESULTS

### Rationale of KAD profiling and software development

With the availability of long-noisy-read and short-accurate-read sequencing technologies, genome sequences nowadays are often constructed by using both long and short whole genome shotgun (WGS) reads. The Illumina sequencing platform is the dominant short-read technology and produces accurate reads with ~0.1% error rate, predominated by single nucleotide substitutions (30). High-depth sequencing data (e.g., 30x or above) and relatively uniformly distributed reads across the genome enable to quantify genome content through k-mer analysis. Specifically, the abundance of a k-mer from short reads should be highly correlated with occurrence of the k-mer in the genome. For most genomes, single-copy k-mers each of which is present once in the genome are dominant among all k-mer sequences (non-redundant k-mers with one or multiple copies) derived from the genome. The mode of sequencing depths of single-copy k-mers (*m*), representing sequencing depth of read data, can be estimated from the spectrum of abundance of read k-mers that are k-mers generated from sequencing reads. For a given read k-mer, the k-mer abundance, or the count per k-mer in reads, is signified by *c*. The occurrence or the copy of the k-mer in a given genome can be estimated by 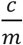. In assembled sequences, the occurrence of the k-mer is signified by *n*. Therefore, 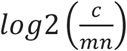 represents the copy number difference between the estimate by reads and the copy of the k-mer in assembled sequences. Because the *n* value per k-mer is 0 for the k-mers that are present in reads but absent in assembled sequences, the formula was adjusted as 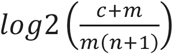, the value of K-mer Abundance Difference (KAD). Using this formula, KADs of k-mers with matching copies indicated by reads and the assembly should be 0 or around 0. If a single-copy k-mer from an assembly is resulted from errors, and no such k-mer is found in reads, the KAD equals −1. Read k-mers missed in the assembly have positive KAD values. In such cases, high-copy k-mers from a genome that are well represented in reads but not in the assembly have high KAD values. Collectively, this simple KAD metric indicates how each k-mer matches with read data in copy number. Therefore, based on the KAD profile of all k-mers together, the quality of an assembly can be assessed using a common standard informed by a read set.

### Base errors in assemblies can be detected through KAD analysis

The use of KAD in error detection was tested by the simulation using an *Escherichia coli* reference genome (4.7 Mb). The KAD calculation requires a genome assembly and short reads generated from the genome. We simulated 50x reads without errors (error-free reads) from the *E. coli* reference genome and 10 sets of genome sequences with 0.1-1% single nucleotide substitution errors (Methods). The KAD value using 25-mer as the k-mer size was determined for each k-mer derived from simulated genomes that contains varying numbers of errors. A k-mer with the KAD value equaling −1 was referred to as an error k-mer. As expected, the numbers of error k-mers increased with error rates of the simulated genomes (**Figure 1A**). We extended simulations using larger genomes from 4 additional species, namely, yeast (*Saccharomyces cerevisiae,* 12.4 Mb), Arabidopsis (*Arabidopsis thaliana*, 121.6 Mb), rice (*Oryza sativa japonica,* 381.3 Mb), and maize (*Zea mays,* 2.2 Gb). For each genome, 50x error-free reads were simulated. Similar to the simulation using the *E. coli* genome, the number of error k-mers detected by KAD analysis in four species showed a linear correlation with error rates (**Figure 1A, Figure S1, Table S1**), which supported the conjecture that the number of error k-mers determined by KAD analysis accurately reflects error rates regardless of genome size and complexity.

**Figure 1.**
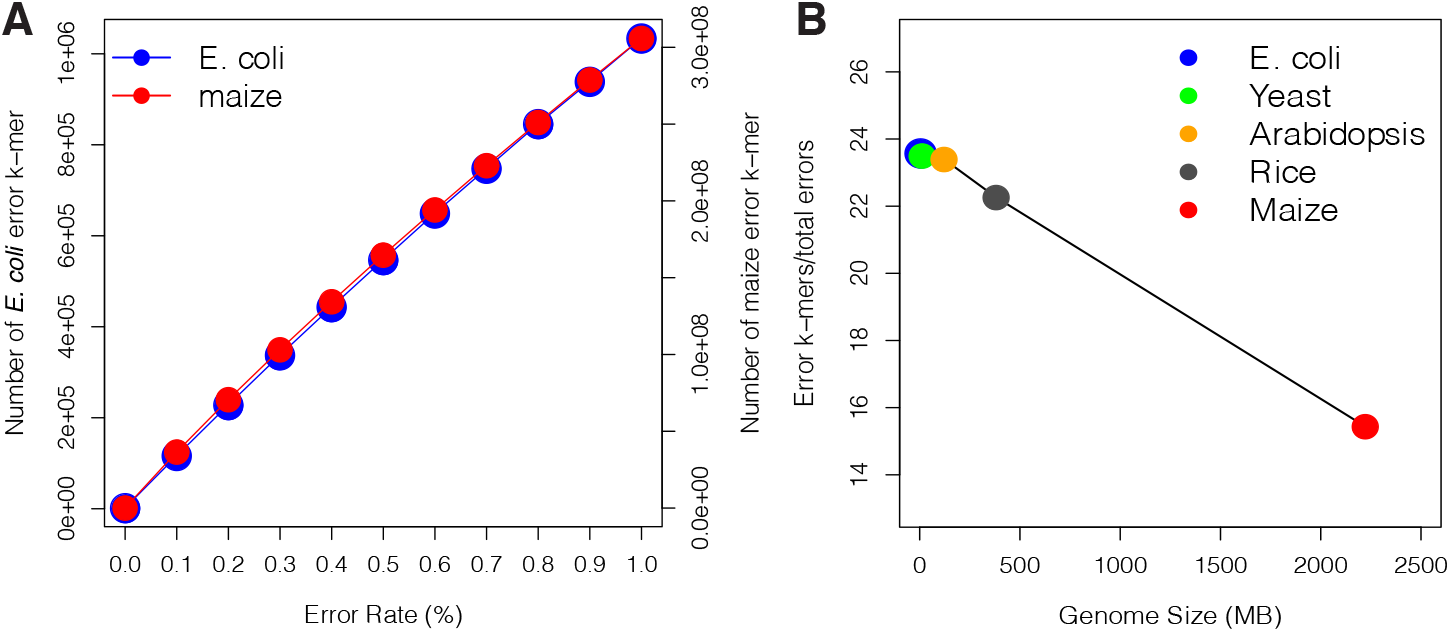
Error detection through KAD analysis. (**A**) Number of error k-mers detected in simulated *E. coli* and maize genomes with different simulated error rates. The left and right y-axes represent the number of error k-mers detected in the simulated *E. coli* and maize genome sequences, respectively. (**B**) Under the 0.5% rate of errors in simulated genomes, ratios of the number of error k-mers to the number of simulated errors in five simulated genomes were plotted versus corresponding genome sizes.

Error k-mers were mapped to simulated genomes and the regions covered by error k-mers were referred to as error regions. The simulated errors located in error regions were considered as errors captured by KAD and the percentage of captured errors was defined as True Capture Rate (TCR). In relatively small genomes (*E. coli*, Yeast, and Arabidopsis) with error rates ranging from 0.1% to 1%, the TCR values were all higher than 99% (**Table 1**), which indicated that KAD analysis can inform almost all errors in these simulated genomes. For a genome with a moderate size (rice), the TCR values reduced to ~95%. The TCR values were further reduced to ~67% for maize that has a large and complicated genome, suggesting that the size and complexity of a genome have impacts on error detection.

**Table 1.**
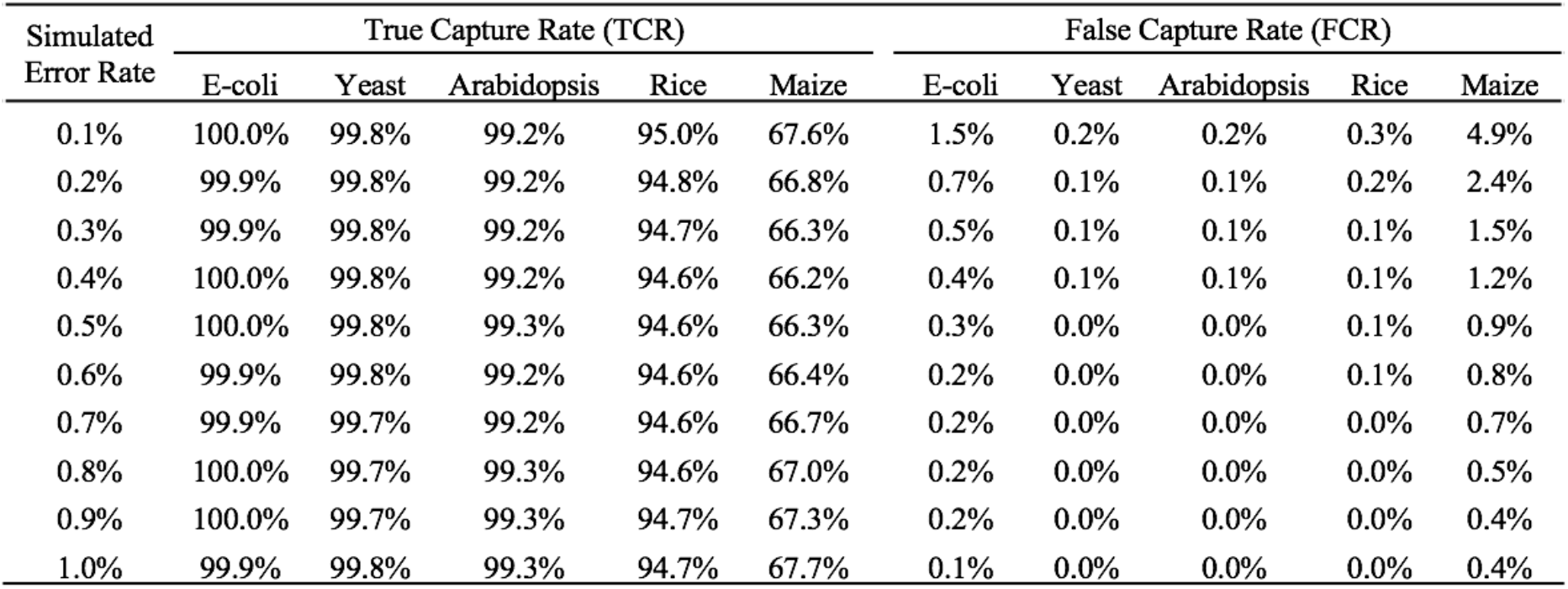
Assessment of error detection via KAD analysis.

| Simulated Error Rate | True Capture Rate (TCR) |  |  |  |  | False Capture Rate (FCR) |  |  |  |  |
| --- | --- | --- | --- | --- | --- | --- | --- | --- | --- | --- |
|  | E-coli | Yeast | Arabidopsis | Rice | Maize | E-coli | Yeast | Arabidopsis | Rice | Maize |
| 0.1% | 100.0% | 99.8% | 99.2% | 95.0% | 67.6% | 1.5% | 0.2% | 0.2% | 0.3% | 4.9% |
| 0.2% | 99.9% | 99.8% | 99.2% | 94.8% | 66.8% | 0.7% | 0.1% | 0.1% | 0.2% | 2.4% |
| 0.3% | 99.9% | 99.8% | 99.2% | 94.7% | 66.3% | 0.5% | 0.1% | 0.1% | 0.1% | 1.5% |
| 0.4% | 100.0% | 99.8% | 99.2% | 94.6% | 66.2% | 0.4% | 0.1% | 0.1% | 0.1% | 1.2% |
| 0.5% | 100.0% | 99.8% | 99.3% | 94.6% | 66.3% | 0.3% | 0.0% | 0.0% | 0.1% | 0.9% |
| 0.6% | 99.9% | 99.8% | 99.2% | 94.6% | 66.4% | 0.2% | 0.0% | 0.0% | 0.1% | 0.8% |
| 0.7% | 99.9% | 99.7% | 99.2% | 94.6% | 66.7% | 0.2% | 0.0% | 0.0% | 0.0% | 0.7% |
| 0.8% | 100.0% | 99.7% | 99.3% | 94.6% | 67.0% | 0.2% | 0.0% | 0.0% | 0.0% | 0.5% |
| 0.9% | 100.0% | 99.7% | 99.3% | 94.7% | 67.3% | 0.2% | 0.0% | 0.0% | 0.0% | 0.4% |
| 1.0% | 99.9% | 99.8% | 99.3% | 94.7% | 67.7% | 0.1% | 0.0% | 0.0% | 0.0% | 0.4% |

We also determined the False Capture Rate (FCR), which is the ratio of error k-mers that do not overlap with simulated errors to total error k-mers. Using data with simulated errors ranging from 0.1% to 1%, all FCR values remained at low levels (<5%) in all simulated genomes (**Table 1**), indicating that KAD analysis accurately pinpointed errors. In addition, in the range of 0.1-1% simulated error rates, the number of error k-mers detected exhibited an approximately linear relationship with the number of errors regardless of the genome size. Thus, we can use the number of error k-mers to estimate the number of errors in a genome assembly. Our simulation data showed that, depending on the error rate and the genome size, the exact conversion ratio of the number of error k-mers to the error rate varied from 14 to 25 when 25-mer was used as the k-mer length (**Figure 1B, Figure S2, Table S2**). In addition, we performed KAD analysis using 31-mer for five genomes with 1% simulated errors, both TCR and FCR retained very high and very low respectively for all genomes except the complex maize genome, for which TCR was improved from 68% to 80% and FCR was reduced from 0.4% to 0.3% (**Table S3**). The TCR and FCR values continued to improve when higher k-mers (37, 43, 49, and 55 mers) were used for maize analyses using the same simulated assembly and read data (**Table S4**). Among them, the 49-mer had the highest TCR (91.9%) and the lowest FCR (0.12%). Collectively, the simulation supported that KAD analysis can detect most errors from the assemblies of small genomes to those of large genomes and provided the extrapolation formula to estimate error rates based on the number of error k-mers detected by KAD analysis.

### KAD analysis detects diverse types of errors

Besides single nucleotide substitution errors, we predicted that KAD analysis can inform other types of errors, such as short insertion and deletion (INDELs), missing sequences or assembly collapses, and contaminated DNA sequences in genome assemblies (31). We grouped k-mers other than error k-mers based on their KAD values into: k-mers with KADs close to 0 (Good), k-mers over-represented in the assembly (OverRep), k-mers under-represented at a low level in the assembly (LowUnderRep), and k-mers under-represented at a high level in the assembly (HighUnderRep) (**Figure 2**). Good k-mers represent k-mers from well assembled and accurate sequences. OverRep k-mers are k-mers with low KAD values (e.g., smaller than −1), representing k-mers showing multiple times in the assembly but are absent or having lower copies indicated by read data. K-mers from redundant sequences in the assembly belong to this group. Under-represented k-mers are k-mers that occur less frequently in the assembly. K-mers derived from incompletely assembled or missing sequences belong to LowUnderRep or HighUnderRep. If a repetitive sequence has a high number of copies and most are missing, the derived k-mers would have a high KAD value (e.g., greater than 2) and therefore would be grouped to HighUnderRep. Note that the KAD ranges specified here are default values used in the KAD scripts to define each k-mer group, and may be redefined in different applications.

**Figure 2.**
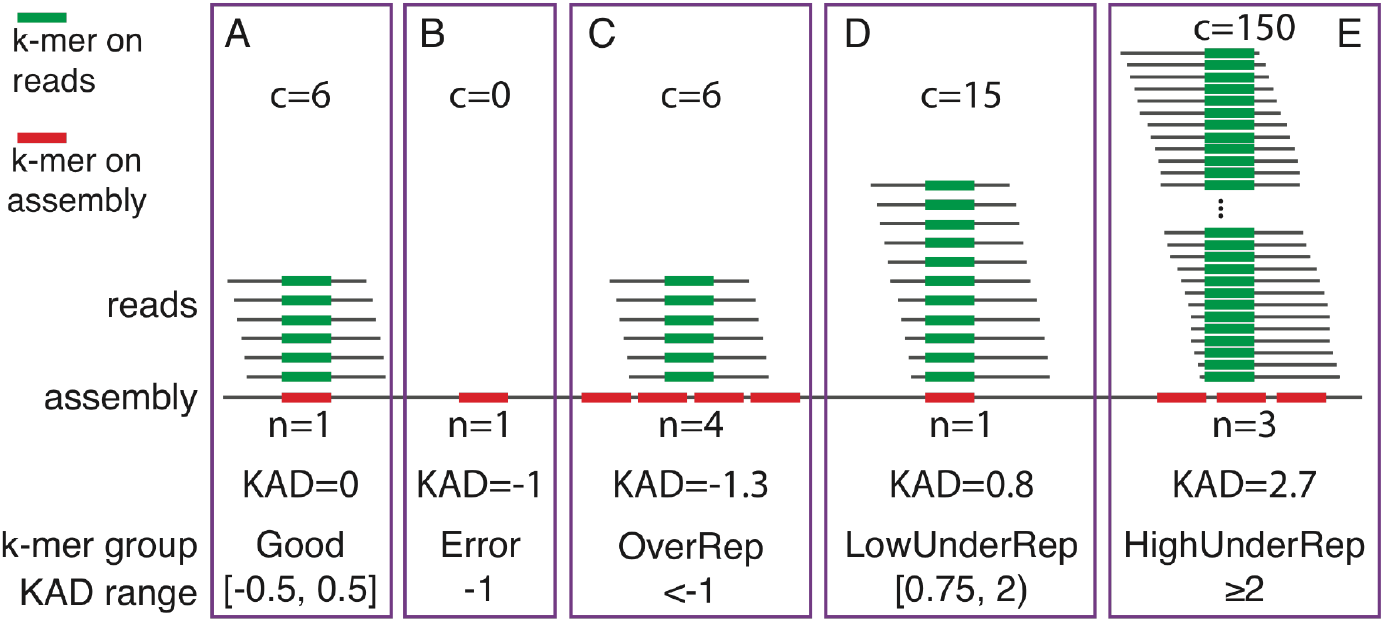
K-mer classification based on KAD values. Examples are provided to illustrate from each k-mer group (**A**) Good, (**B**) Error, (**C**) OverRep whose k-mers are over-represented in the assembly, (**D**) LowUnderRedp whose k-mers are under-represented at a low level in the assembly, and (**E**) HighUnderRep whose k-mers are under-represented at a high level in the assembly. In these examples, 6 reads are the mode of k-mer abundances (*m*=6). The *c* and *n* values are abundances of k-mer in reads and copy numbers in the assembly, respectively. In each purple box, green bars highlight a k-mer in reads and red bar(s) indicate the occurrence of the k-mer in the assembly. The KAD ranges specified are cutoff values that define each k-mer group.

We next examined how the KAD-based grouping strategy detects various assembly errors through simulation of multiple types of errors using the *E. coli* genome. We separately simulated four types of errors: (I) single nucleotide substitution ranging from 1% to 10% (II) short INDELs (less than 10 bp) ranging from 1% to 10% (III) long sequence redundancy (insertion between 100 bp and 1000 bp) and (IV) sequence deletion (deletion between 100 bp and 1000 bp) with the number from 50 to 500. KAD analysis was performed on each of these simulated genomes along with 50x error-free reads (**Table S5**). For the types I and II, while the error rates increase, the number of Good k-mers decreases and the number of Error k-mers increases (**Figure 3A, 3B**). Because of the presence of error sequences, corresponding correct sequences are under-represented in the assembly, resulting in the increase of k-mers in the group of “lowUnderRep”. Note that, owing to a higher frequency of multiple errors in an error k-mer when the error rate in the simulated genomes is greater than 2%, the linear relationship between the number of error k-mers and error rates was not maintained (**Figure 3A, 3B**). Long sequence errors, redundancy and deletion, in the assembly resulted in abundant OverRep k-mers and LowUnderRep k-mers, respectively. But both of them had a few error k-mers compared to single nucleotide substitutions and short INDELs (**Figure 3C, 3D**, **Table S5**). These results indicated that KAD analysis was able to separate errors due to sequence redundancy or missing assemblies. At the same time, the KAD analysis can detect errors caused by single nucleotide substitutions and short INDELs but cannot distinguish these two error types.

**Figure 3.**
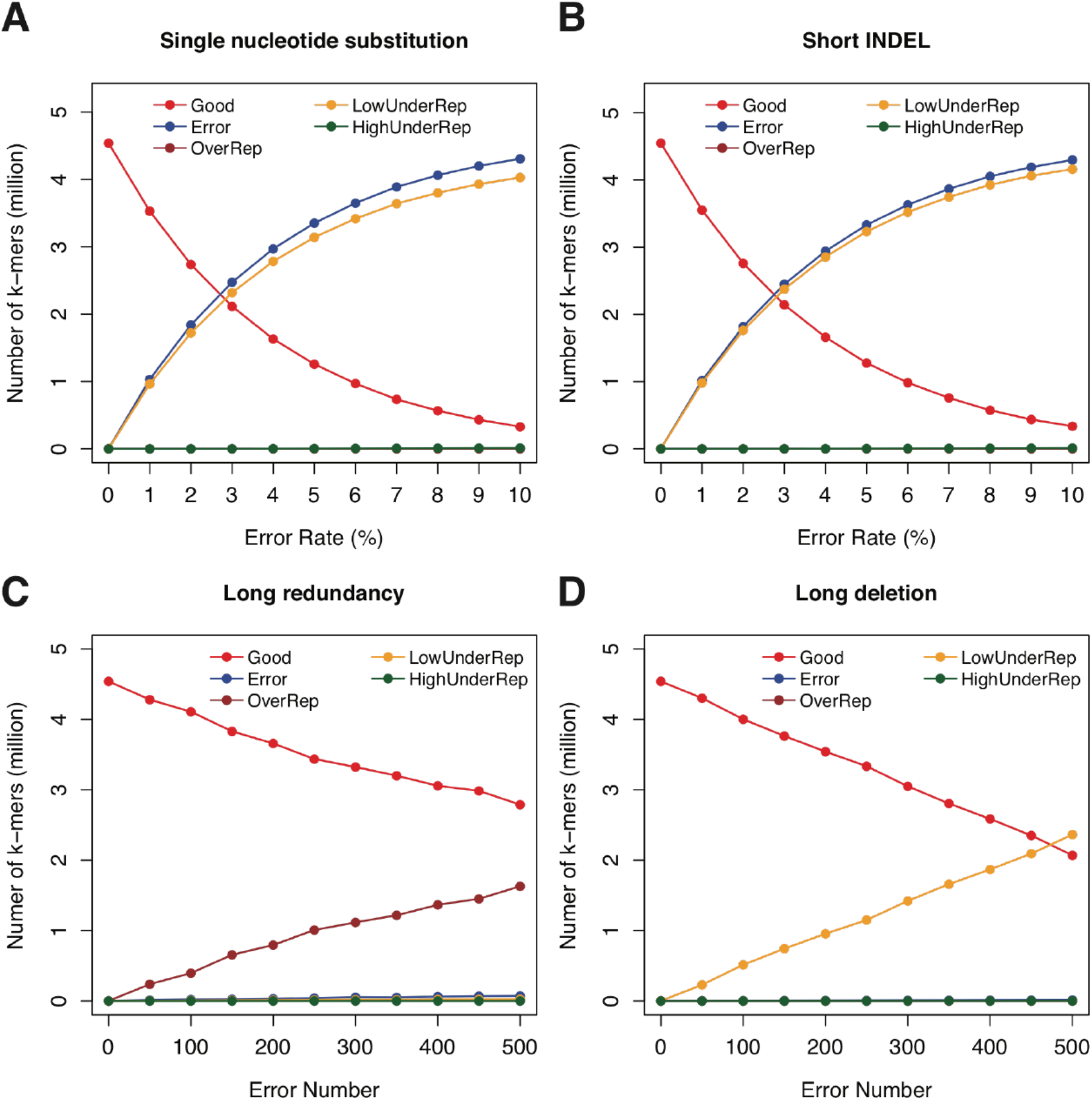
KAD analysis of multiple error types simulated on the *E. coli* genome. KAD results of simulated genome sequences with different types of errors: (**A**) single nucleotide substitutions, (**B**) short INDELs, (**C**) long sequence redundancy, and (**D**) long sequence deletion. In each plot, colors of lines represent different groups of k-mers detected by KAD.

### The impacts of depths and errors of sequencing reads on KAD analysis

KAD analysis quantitatively compares k-mer abundance in sequencing reads and in the examine the impact of sequencing depth on KAD analysis, error-free reads were separately simulated with read depths from 10x to 100x in *E. coli* genome. Using various depths of sequencing read data, KAD analysis was performed on the previously simulated genome sequences with 1% to 10% single nucleotide substitution errors. As a result, the accuracy of error detection using KAD analysis, represented by TCR, was >99.9% for simulated *E. coli* genomes when sequencing depths were higher than 20x (**Figure 4A**). However, when sequencing depth is lower than 40x, false error capture rates (FCR) were high for the simulated genomes with low error rates (**Figure 4B**). Similar results were observed for simulations with other three larger genomes, namely, yeast, Arabidopsis, and rice, with 0.1% to 1% single nucleotide substitution (**Data S1**). This simulation result indicated that at least 40x read depth is required for accurate detection of errors through KAD analysis.

**Figure 4.**
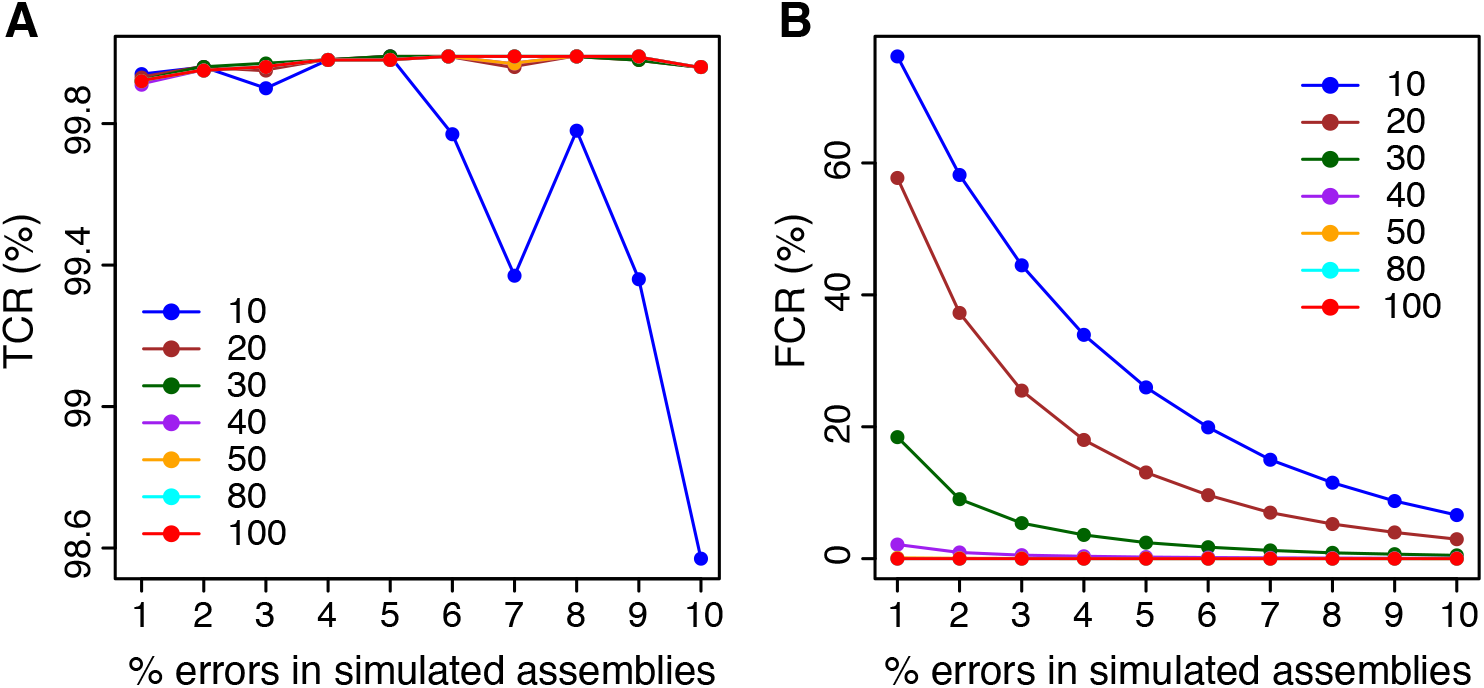
Impacts of read depths on error detection in simulated genomes. Various random errors of single nucleotide substitution on the *E. coli* reference genome were simulated, resulting in simulated *E. coli* genomes with errors ranging from 1% to 10% that are shown on x-axes. KAD analyses were performed to detect errors using different depths of error-free read data whose depths range from 10x to 100x coverages. Both TCR (**A**) and FCR (**B**) of error detection *via* KAD were determined, and plotted versus percentages of simulated errors in the simulated genomes. Each curve represents a certain depth of read data used for KAD analyses. Depths of read data, ranging from 10 to 100x, are color coded and labeled within each plot.

All simulations so far used error-free reads. To examine the impact of read errors on KAD analysis, 50x reads with error rates of single nucleotide substitutions from 0.1% to 0.5% and from 1% to 5% were simulated from the *E. coli* reference genome. KAD analysis was then performed using these error-bearing reads and the simulated *E. coli* genomes with 1% to 10% single nucleotide substitutions. The numbers of error k-mers detected by KAD were robust using reads with ≤2% error rates (**Figure 5A**). The TCR values stayed above 99.9% for all error rates of reads from 0.1% to 5% (**Figure 5B**). However, where error rates of reads were higher than 3%, high levels of FCR were observed (**Figure 5C**). This simulation result showed that, as long as read errors are not higher than 1%, the impacts of read errors on error detection of assembled sequences through KAD analysis is trivial.

**Figure 5.**
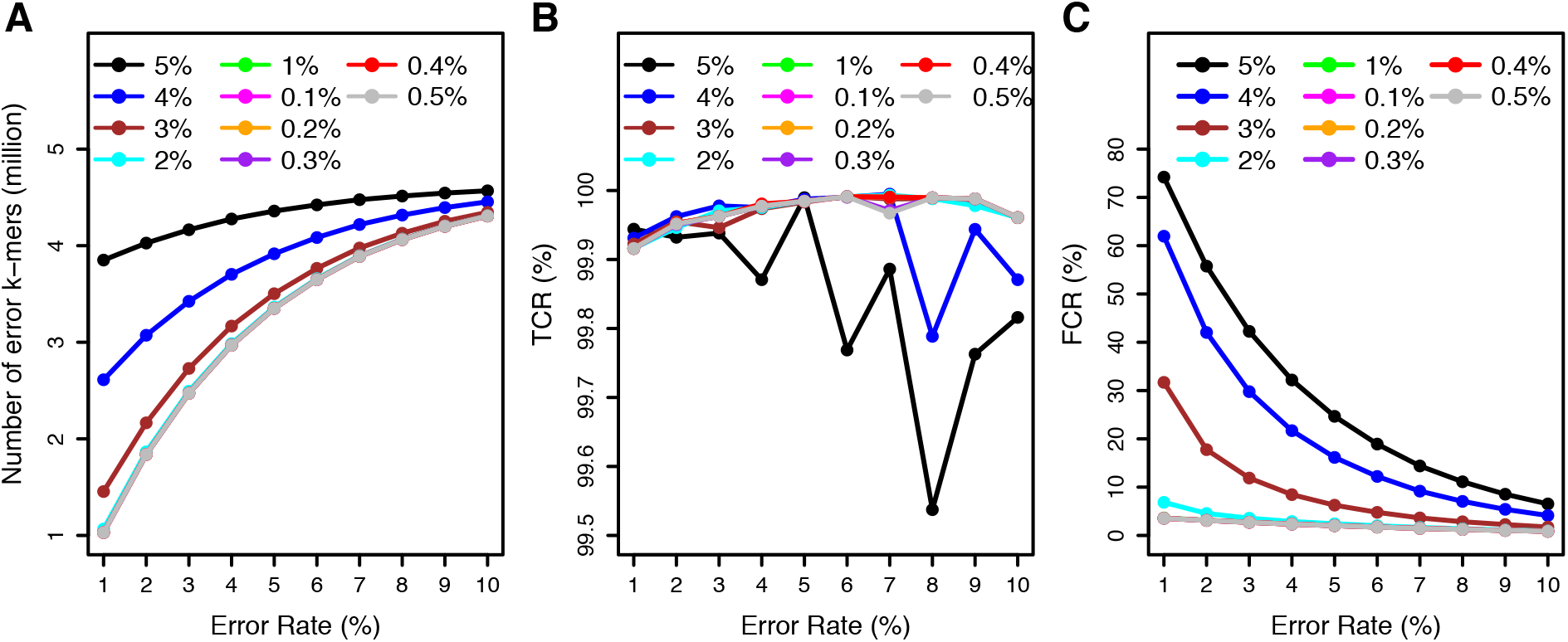
Impacts of read errors on error detection. The simulated *E. coli* genome sequences contain single nucleotide substitution errors ranging from 1% to 10%. KAD analyses were performed to detect errors on these simulated genomes using error-bearing reads ranging from 0.1% to 0.5% and from 1% to 5%. Numbers of error k-mers detected (**A**), TCR of error detection (**B**), and FCR of error detection (**C**) were plotted versus error rates in simulated genome sequences. Each curve represents a certain level of errors in reads used for KAD analysis. Different levels of errors in reads are color-coded and labeled in each plot.

### Assessing a bacterial genome assembly via KAD analysis

We previously produced a genome assembly of a *Xanthomonas vasicola* pv. *vasculorum* (Xvv) isolate Xvv1601 that was isolated from a Kansas corn adult leaf showing the symptoms characteristic of bacterial leaf streak in 2016 (32). PacBio long reads were used for the genome assembly with the assembler Canu (26). The resulting assembly with the software Canu, referred to as assembly canu, was then polished with raw PacBio reads and Illumina reads, followed by the circularization by removing overlapping ends, resulting in a final assembly (assembly final, Genbank accession CP025272.1). The final assembly consists of 4,956,923 bp. Comparison between assemblies canu and final found 142 polymorphisms that were all one-base INDELs.

KAD analysis was performed on two versions of assemblies with trimmed Illumina 2×300bp paired-end reads. The spectrum of k-mer abundance showed that the sequencing depth of Illumina reads is 199 (**Figure 6A**). KAD profiling of both assemblies canu and final showed KAD values of most k-mers are around 0, indicating that the overall base quality of both assemblies is high (**Figure 6B**). In assembly canu, a small peak of error k-mers (N=2,649) is detected, suggesting that base errors were retained in assembly canu. These error k-mers covered 141 error regions on assembly canu. All error regions were not larger than 51 bp. Strikingly, all 142 INDELs between the two versions of assemblies were located within these small error regions and each error region contained one INDEL polymorphism except one that contained two. KAD analysis of assembly canu also showed that some k-mers are under-represented in the assembly (**Figure 6C**). In the final assembly, no error k-mers were found but under-represented (both LowUnderRep and HighUnderRep) k-mers largely retained (**Figure 6D**). Aligning these under-represented k-mers to Genbank databases found that these k-mers were not derived from bacterial genomes. Instead, they were from the PhiX phage DNA that was used as controls for Illumina sequencing or DNA sequences of organisms other than *Xanthomonas*. Therefore, these sequences under-represented in the assembly were likely generated from DNA contamination during Illumina sequencing or library preparation.

**Figure 6.**
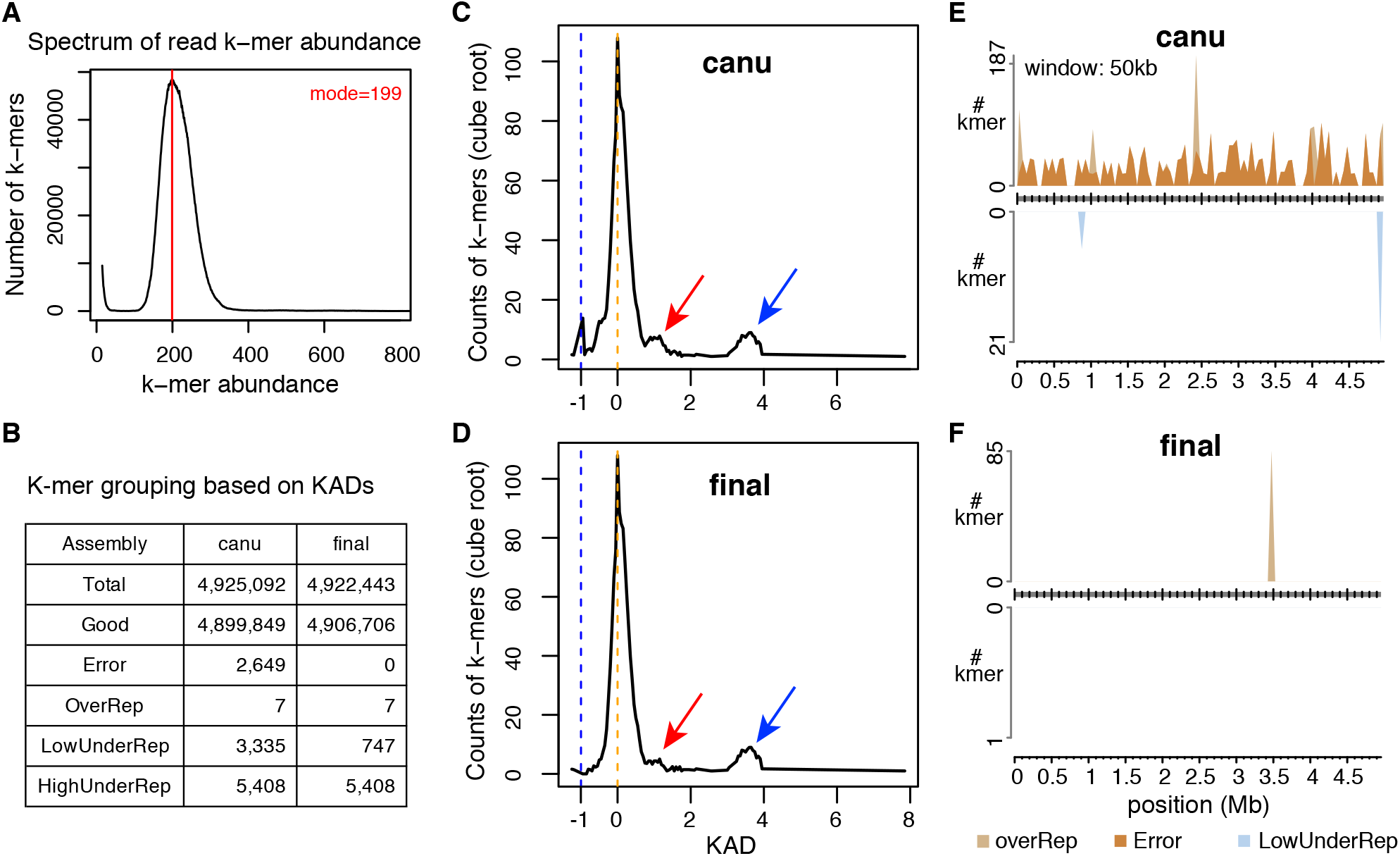
KAD analysis of assemblies of a bacterial genome. (**A**) Spectrum of abundance per k-mer profiled with trimmed Illumina reads. The spectrum determined the sequencing depth of Illumina data. (**B**) The summary table of numbers of k-mers in each k-mer group. Both k-mer counts from the unpolished assembly (canu) and the polished assembly (final) are shown. (**C**) KAD profiling of assembly canu shows that a strong peak is at around the KAD value of 0 (orange dash line), which represents correct k-mers (referred to as “Good”); a small peak at the KAD value of −1 (blue dash lines), which represents a group of error k-mers (referred to as “Error”). Red arrow points at a small bump representing a low-level of under-represented k-mers in the assembly, referred to as “LowUnderRep”. Blue arrow points at a bump representing a high-level of under-represented k-mers in the assembly, referred to as “HighUnderRep”. (**D**) The KAD profile of the “final” assembly. The peak representing error k-mers is disappeared. However, small bumps (red and blue arrows) remain, which contains k-mers from contaminated DNA during Illumina sequencing. (**E**) Grouped k-mers were mapped to assembly canu. All highUnderRep k-mers were not mapped. (**F**) Grouped k-mers were mapped to assembly final. Seven OverRep k-mers were repeatedly mapped to a 15-copy tandem “ATTCGGG” 7-bp repeat. In both **E** and **F**, and the number of k-mers per 50 kb in each group was determined and plotted versus the position of the 50-kb window in the assembly. A k-mer was counted multiple times if it was mapped to different locations.

To understand distributions of k-mers of different groups, k-mers were mapped to both canu and final assemblies, which required perfect matches but allowed multiple mapping locations. A KAD landscape plot displaying distributions of grouped k-mers from assembly canu showed that error k-mers were spread along the whole assembled genome and a few OverRep and LowUnderRep k-mers were mapped (**Figure 6E**). The KAD landscape plot of assembly final located the OverRep k-mers at a region that has 15-copy “ATTCGGG” 7-bp tandem repeats (**Figure 6F**). Collectively, KAD analysis indicated that the final assembly of Xvv1601 (CP025272.1) is a finished genome assembly with a very high base quality.

We randomly sampled reads from original high-depth (~200x) reads to the coverages from 90x to 20x and performed KAD analysis using each down-sampled read dataset. When the read depth was 40x or above, at least 98% error k-mers originally identified in assembly canu were repeatedly identified with the total number of error k-mers highly close, and 0 error k-mers were found in the final assembly (**Figure S3**). However, both error k-mers in assemblies canu and final were slightly increased when the read depth was 30x, but were dramatically increased when the read depth was 20x (**Figure S3**). The result suggested 40x is the minimum read depth for accurate KAD analysis, which agreed with previously simulation results.

### KAD analysis to improve the genome assembly of a fungal wheat blast isolate

We previously used PacBio long reads to assemble a near-finished genome assembly of fungal field isolate B71 that causes wheat blast diseases (28). The B71Ref1 assembly consists of seven core-chromosomes, five contigs that are from a supernumerary mini-chromosome, and a mitochondrial sequence (28). To improve the assembly, we generated Nanopore long reads (**Figure 7A, Figure S4**) and performed a *de novo* assembly using only Nanopore dada, resulting in 12 contigs. Based on the alignment of polished contigs (ONTv0.14, Methods) with the previously B71Ref1 assembly, we reorganized these contigs into chromosomes and corrected a misassembly between chromosome 6 and a verified mini-chromosome sequence (**Figure S5**). The updated Nanopore assembly contains seven core-chromosomes, a mini-chromosome, and a mitochondrial sequence. We also updated the previous PacBio assembly with additional polishing steps (PBRef1.3). To compare ONTv0.14 and PBRef1.3, we performed KAD profiling on both assemblies and aligned the two assemblies using Nucmer (27). The alignment of each core-chromosome indicated that the two assemblies have a very low level of dissimilarity (**Figure 7B, Figure S6**). KAD analysis enabled the determination of errors and their distribution along each chromosome. Combining both alignment results and error profiling indicated by KAD analysis, we selected assembled sequences from either assembly that contains more complete sequences or fewer errors (**Table S6**). The combined assembly (B71Ref1.5) took advantage of both Nanopore and Pacbio long read technologies. KAD profiling of B71Ref1.5 indicated that the combined assembly carries ~2,568 base errors as estimated by the number of error k-mers, i.e., 99.995% accuracy (**Table S7**).

**Figure 7.**
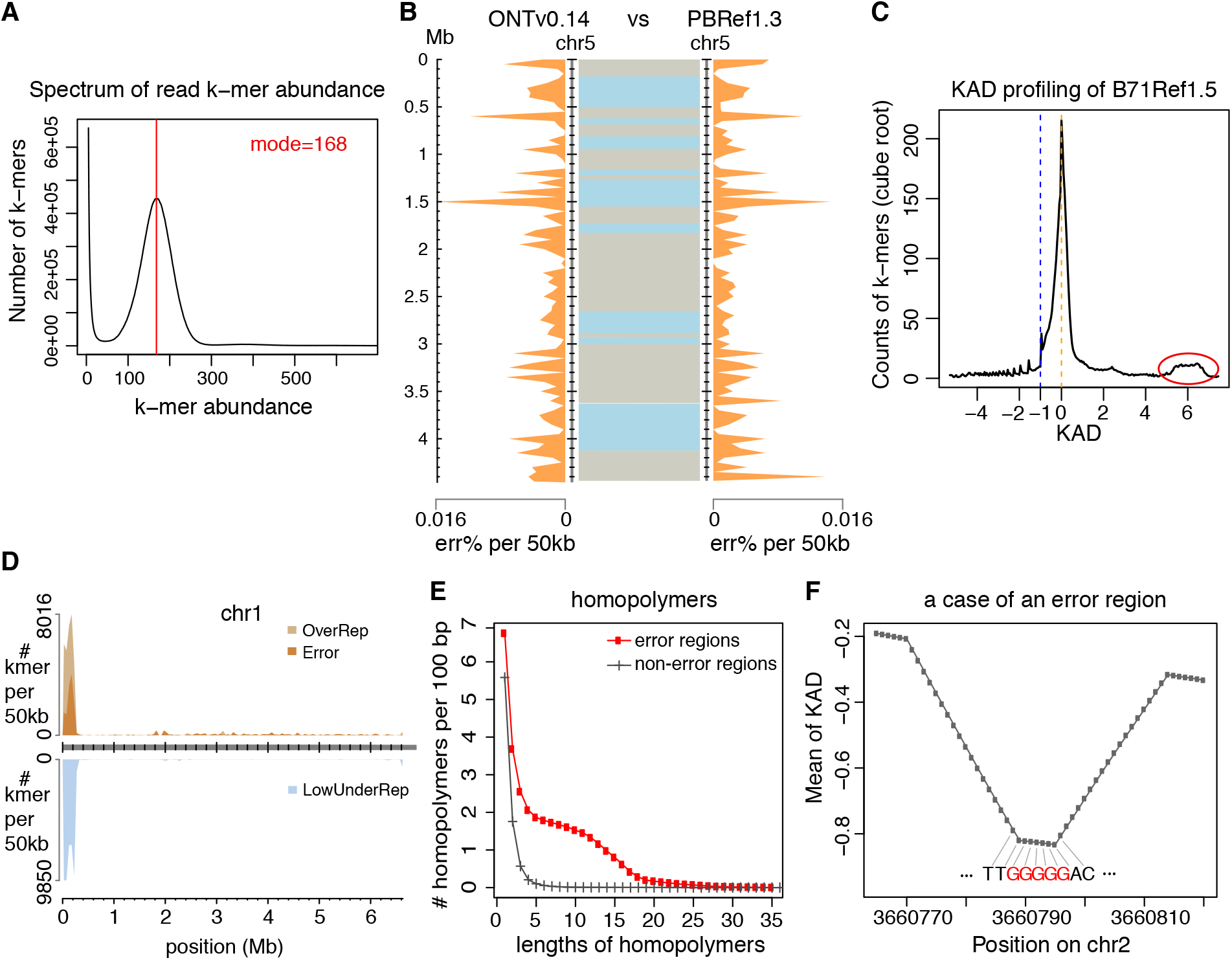
KAD analysis identified errors in the fungal genome assembly. (**A**) Spectrum of abundance per k-mer profiled with trimmed Illumina reads, which was used to determine the sequencing depth. (**B**) Comparison of chromosome 5 between the assembly with Nanopore long reads (ONTv0.14) and the assembly with PacBio long reads (PBRef1.3). Distributions of percentages of estimated base errors (err%) per 50 kb are plotted separately for the two assemblies along chromosome 5 (orange shades). In between two assemblies, alignments of the same chromosomes with NUCmer are displayed with two colors to indicate neighboring alignments. (**C**) B71Ref1.5 is the merged assembly combining both ONTv0.14 and PBRef1.3. KAD profiling was performed using this merged assembly and trimmed Illumina reads. The red circle highlights a bump that is largely contributed by k-mers from the mitochondria whose genome was in high number of copies in each cell. (**D**) Error k-mers and under-represented k-mers identified using the B71Ref1.5 assembly were mapped to the B71Ref1.5 assembly. Each k-mer was allowed to map to up to 100 locations. The number of k-mers in each group per 50 kb was determined and plotted versus the position of the 50-kb window in the assembly. (**E**) Counts of homopolymers of different lengths per 100 bp were plotted versus lengths of homopolymers. The red curve represents counts on error regions and the gray curve represents counts on non-error regions. (**F**) After k-mers were aligned to the assembly, the mean KAD value per position was calculated by averaging KAD values of all k-mers aligned to the position. This example shows the mean KAD values per position at a small region on chromosome 2. The DNA sequence is from the assembly. Each base matches the linked position. The lowest mean of KAD points at a G homopolymer tract (red).

More than 99% error k-mers were uniquely and perfectly mapped to assembled sequences. The KAD landscape plot showed that error k-mers were spread along the whole genome but obviously not randomly distributed (**Figure S7**). About 66% error k-mers were located in the repetitive regions that consist of only 13% of the genome. K-mers with KAD values greater than 5 were largely derived from the mitochondrion that had a high number of copies each cell (**Figure 7C**). Most k-mers from the highly repetitive ribosome DNA (rDNA) at the beginning of chromosome 1 have high KAD values (around 2), indicative of incomplete assembly. The rDNA cluster region also carries a higher level of errors (~0.4%) (**Figure 7D**). We defined error regions where error k-mers were mapped. Sequence analysis of these error regions revealed that the homopolymers with the length between 4 and 16 are highly enriched as compared to their frequencies in the genomic regions with no error k-mers mapped (**Figure 7E**). We also determined the means of KAD values per genomic position by averaging KADs values of k-mers mapped to the position. At small regions with negative continuous KAD values, we frequently observed the positions with the lowest means of KADs pointed at homopolymer tracts (as an example shown in **Figure 7F**). Here we scrutinized a Nanopore and Pacbio combined wheat blast fungal assembly, we showed that the combined assembly B71Ref1.5 is a “finished” assembly using the community standard (33) but is still incomplete at highly repetitive regions and contains a low level of base errors, particularly in repetitive sequences.

## DISCUSSION

Remarkable progress has been made to advance genome assemblies in recent decades, including cost-efficient and accurate high-throughput short-read sequencing, long-read single molecule sequencing, and improved assembly and error correction algorithms (34, 35). The evaluation of multiple versions of assemblies using different assembly algorithms and various assembly procedures is critical for optimization of genome assemblies. It is also important to assess the final assembly products for information on errors and incompleteness. Here we developed a simple but effective method for genome evaluation based on the quantitative comparison of k-mer abundances between accurate short-read sequencing data and the assembly sequences. The method, referred to as KAD, depicts how sequence contents from high-depth short-reads are represented in each examined genome assembly, identifies the regions in the assembly where base errors occur, estimates the overall error rate of an assembly, and finds the regions containing potential sequence redundancy or incompleteness. Additionally, problems with short-read data, if they exist, could be detected. The KAD method has been implemented and the scripts are freely accessible (https://github.com/liu3zhenlab/kad), which would be useful for genome assembly projects of a wide range of species from small bacterial genomes to large eukaryotic genomes.

KAD first analyzes high-depth (at least 40x) Illumina data to determine the sequencing depth, or the coverage of the genome. This analysis assumes that k-mers having a single copy in the genome are most abundant among all k-mers generated from the genome, which is true for genomes of most species if the k-mer length is sufficiently long (e.g., 25 nt). Caution is needed when analyzing polyploidy genomes such as wheat or heterozygous diploids. Based on that assumption, the k-mer abundance that occurs most frequently (the mode of k-mer abundances) should represent the sequencing depth of single-copy k-mers if the reads contain no sequencing errors. Reads with sequencing errors, even at a low frequency of sequencing errors, generate a very high number of k-mers with small abundance (e.g., 1-3), and can slightly reduce k-mer abundance of each correct k-mer. Such reduction is negligible as it is very small. We thus use the mode of k-mer abundances (the most frequent k-mer abundance that is not small) as the estimate of the sequencing depth, which is an important parameter in the formula to calculate KADs.

For a complete assembly with no errors and unbiased sequencing data with no contamination, the KAD profile would be ideal in that KADs of all k-mers would be around 0. In reality, problematic k-mers with KADs distant from 0 can be found. We categorized problematic k-mers into four groups: over-represented (OverRep), errors (Error), low-level under-represented (LowUnderRep) and high-level under-represented (HighUnderRep). The group of “Error” is well defined. KAD values of all k-mers of this group are −1, which indicates that no such k-mers are produced from sequencing reads but they are present in the assembly. Importantly, the number of k-mers in this group (error k-mers) reflects the sequence error rate of the assembly. Our simulation data indicate that the assembly error rate can be estimated using the number of error k-mers. This estimation is particularly useful nowadays because many genome assemblies retain a number of errors from noisy long reads that are used for assemblies (7). Importantly, most of these errors create new sequences that are not present in the actual genome sequences. Therefore, most error k-mers can be unambiguously mapped to error regions in the assembled sequences even though error k-mers are located in highly repetitive regions. This provides a unique approach to identify errors in repetitive sequences that are error-prone. The other three groups are arbitrarily categorized by defining the range of KADs. KAD thresholds to define these categories are adjustable but the defaults worked well for all the genomes that we tested.

The over-represented group includes k-mers with low KAD values (e.g., smaller than −1), representing k-mers occurring multiple times in the assembly, but are absent or represented less frequently in reads than expected. Mostly likely, these k-mers are derived from sequence redundancy in the assembly, or regions containing systematic errors across multiple locations in the assemblies, or sequences in the genome that are biasedly under-represented in reads. Biased under-representation in reads could occur at extremely highly GC- or AT-rich regions (25). The under-represented groups include k-mers with high positive KAD values (e.g., greater than 0.75), representing k-mers showing fewer copies in the assembly than indicated by reads. For example, k-mers from a region incorrectly collapsed due to tandem repeats would be categorized in these groups. The higher the KAD values, the higher the level of potential assembly incompleteness. However, when k-mers are derived from high-copy organelles or plasmids, the high KAD values reflect the fact that high copies of the sequence are present but only one is probably assembled. In addition, when k-mers are from contamination in short-reads, their KAD values could be high. When mapping over- and under-represented k-mers to the assembly, the issue of multiple mapping is frequently encountered. KAD scripts allow users to tune the parameter of the maximum number of locations, which enables the examination of k-mers that are located at multiple genomic regions. However, users need to bear this in mind that the over- or under-represented signal from a region could be repeatedly shown at multiple locations.

In this study, for most analyses, we chose 25-mer as the k-mer size because it is an optimal size in the genome assembler ALLPATHS-LG for analyzing k-mer abundance spectrum (36) and we have successfully applied it to examine repetitive sequences of maize that has a large complex genome (21). A shorter k-mer size can have a higher resolution to pinpoint error regions but compromises the uniqueness of each k-mer in the assembled genome sequences. In our simulation result, we showed that 25-mer has high power and accuracy for detecting base error in assembled genomes with small or moderate sizes. We also showed that large complex genomes like maize need longer k-mers for improving the uniqueness of k-mers in the genomes. But error regions identified using longer k-mers are wider, more likely cover multiple errors, and the analysis requires more computation resources. In addition, sequencing reads used in KAD analysis are not error-free. The longer the k-mer size, the higher the likelihood that a k-mer from reads would carry errors (37). Therefore, slightly higher depth than 40x would help ensure reliable error detection when a longer k-mer length (e.g., 49-mer) is used.

The development of the KAD analysis was inspired by a motivation to generate a high-quality genome assembly and to develop a method to compare different assemblies for nomination of the final release. With the KAD bioinformatics pipeline, we can quantify the overall sequence error rate and locate errors. Existing error correction algorithms can use the information provided by KAD for targeted error correction, which should reduce false correction. KAD can also help to determine whether one or multiple rounds of polishing were sufficient to meet the convergence criteria, for example, on the basis of the diagnostic plots including KAD error profiles and KAD landscape plots. In the future, new error correction approaches, particularly approaches that are not based on read alignments, could be developed along with KAD for precise error correction in both non-repetitive and repetitive sequences.

## ACKNOWLEDGEMENTS

We thank funding support from the USDA’s National Institute of Food and Agriculture (award no. 2018-67013-28511) and the US National Science Foundation (awards No. 1741090). This is the contribution number 20-209-J from the Kansas Agricultural Experiment Station.

## DATA AVAILABILITY

Bacterial sequencing raw data and genome assembly are available in the NCBI BioProject under the accession number of PRJNA421835. Fungal Nanopore sequencing raw data and genome assembly are available in the NCBI BioProject under the accession number of PRJNA355407.

## CODE AVAILABILITY

KAD codes are available at Github (https://github.com/liu3zhenlab/KAD). Custom scripts used in the manuscript are available at Github (https://github.com/liu3zhenlab/manuscripts) or upon request.

