## Supplementary material for "Estimating Assembly Base Errors Using K-mer Abundance Difference (KAD) Between Short Reads and Genome Assembled Sequences": Figure S, Table S

### 1. SUPPLEMENTARY FIGURES

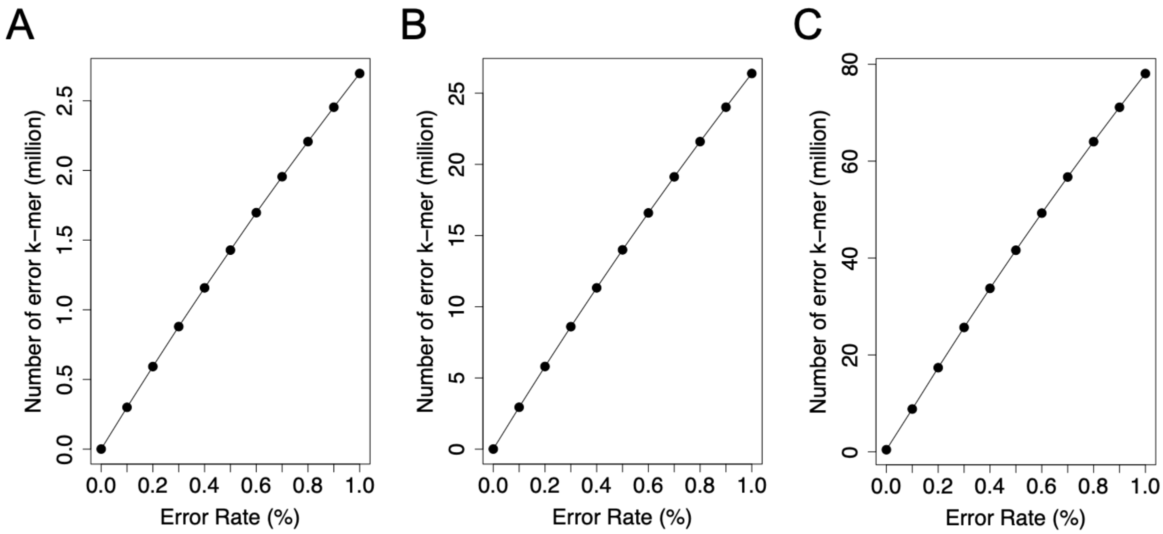


**Figure S1. Error detection of simulated genomes with different error rates.** Numbers of error k-mers detected in simulated genomes were plotted versus rates of errors simulated using the reference genomes from three species: (**A**) *S. cerevisiae*, (**B**) *A. thaliana*, and (**C**) *O. sativa japonica*.

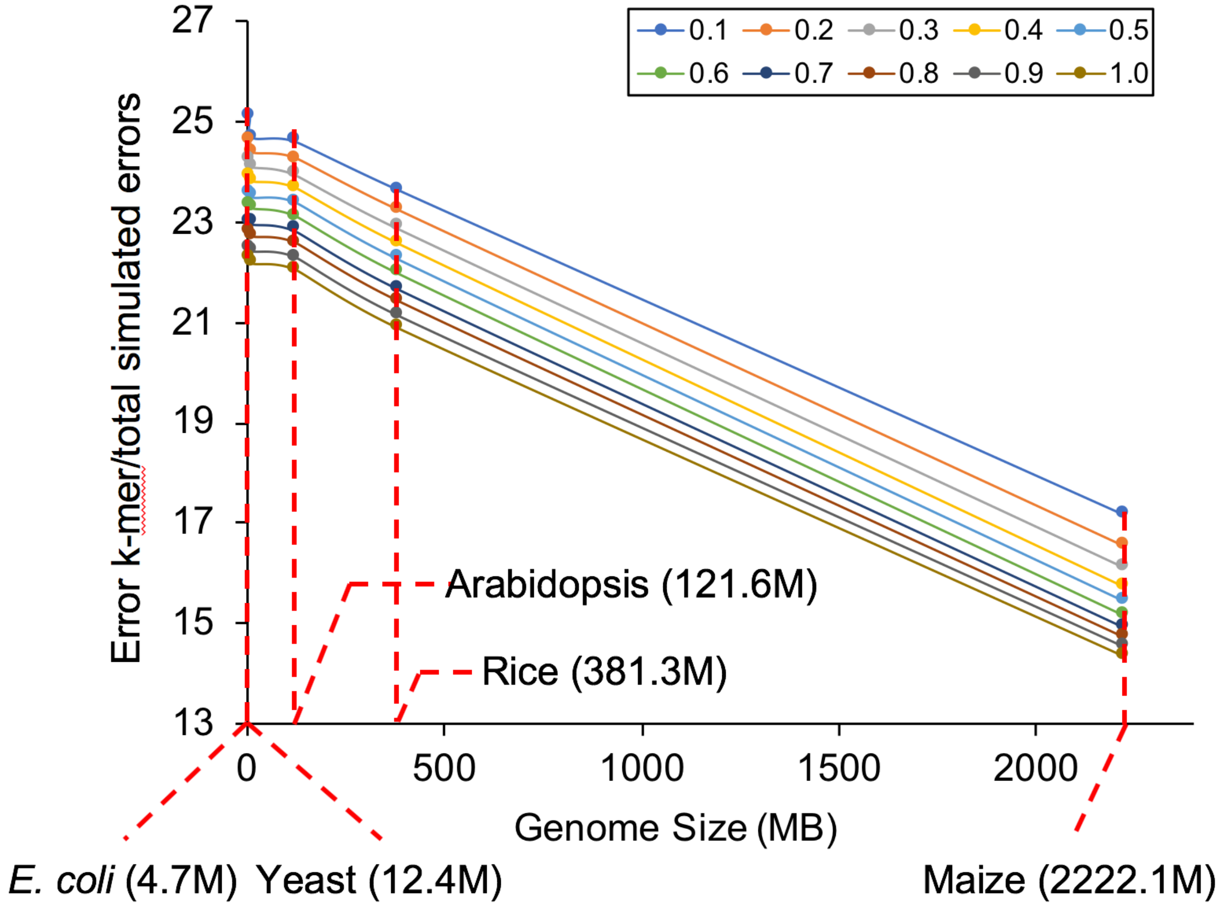


**Figure S2. Ratios of the number of error k-mers to total errors.**
The x-axis represents genome sizes (MB) and the y-axis represents the ratio of the number of error k-mers to the number of total simulated errors. Colors of lines represent simulated error rates in simulated genomes from five species.

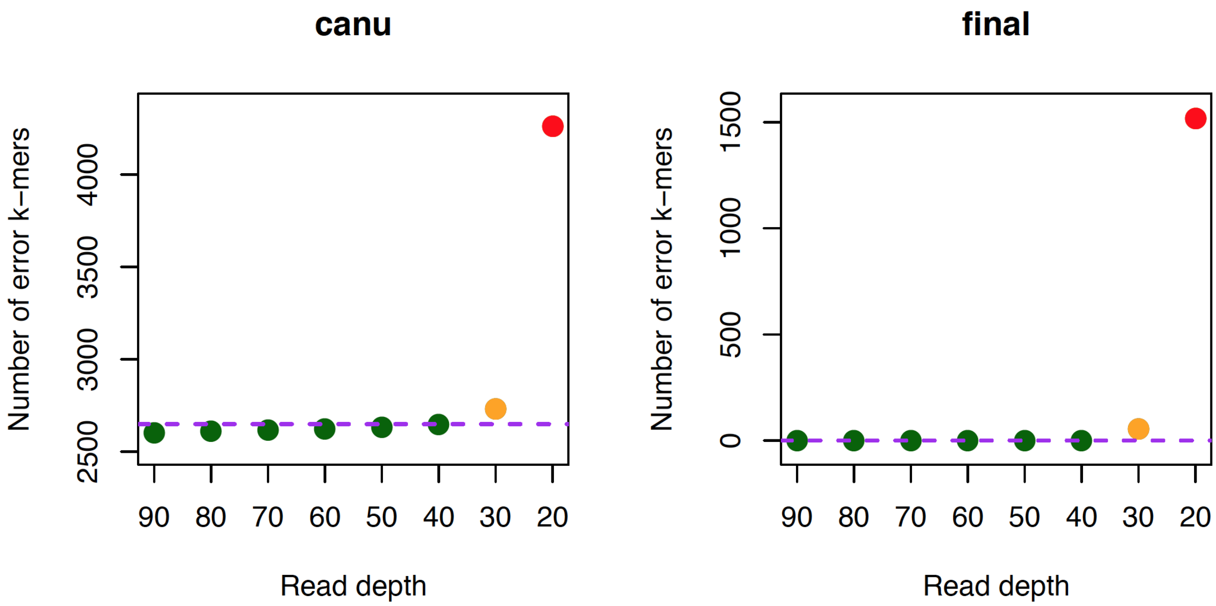


**Figure S3. Number of error k-mers identified with different read depths.** Reads were randomly sampled from ~200x Illumina Xvv1601 paired-end read data to different levels of depths ranging from 90x to 20x. KAD analysis was performed using each down-sampled dataset on both canu and final Xv1601 assemblies. Purple dash lines stand for the numbers of error k-mers identified with the original read depth. Green points represent data points that are very close to the error k-mers identified with original read depths; orange points represent the ones that slightly higher than the error k-mers originally identified; and red points represent the ones that dramatically higher than the error k-mers originally identified.

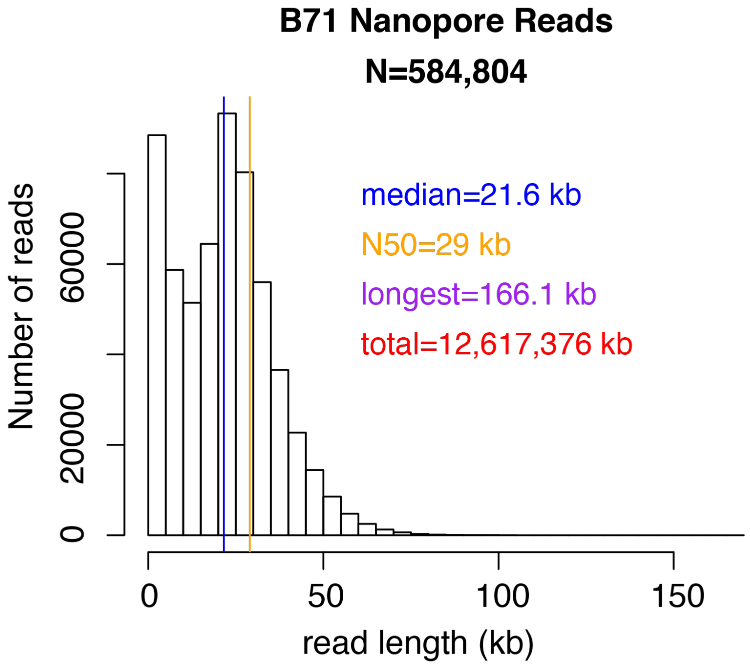


**Figure S4. Histogram of Nanopore read lengths.**
The median and N50 are indicated by blue and red vertical lines, respectively.

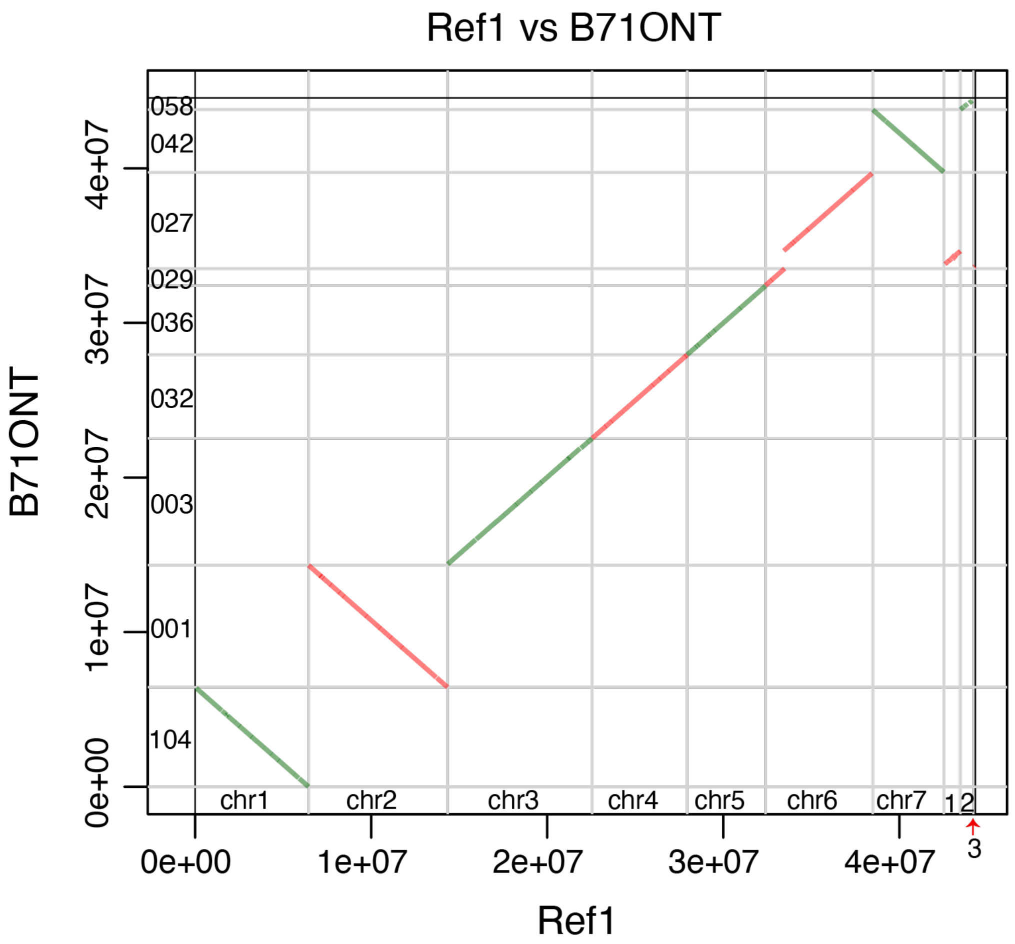


**Figure S5. Dotplot between Nanopore draft assembly and B71Ref1 (Ref1).** Each alignment requires at least 50 kb matches with at least 98% identity. Sequences 1, 2, 3 of Ref1 represent scaf1, scaf2, and scaf3, respectively. Numbers inside y-axis coordinates are contig IDs (last three digits of the original contig IDs in the format of tig00000xxx) of the Nanopore draft assembly. The scaf1 from Ref1 validated from the mini-chromosome was mis-assembled in the contig of “027” of the Nanopore draft assembly.

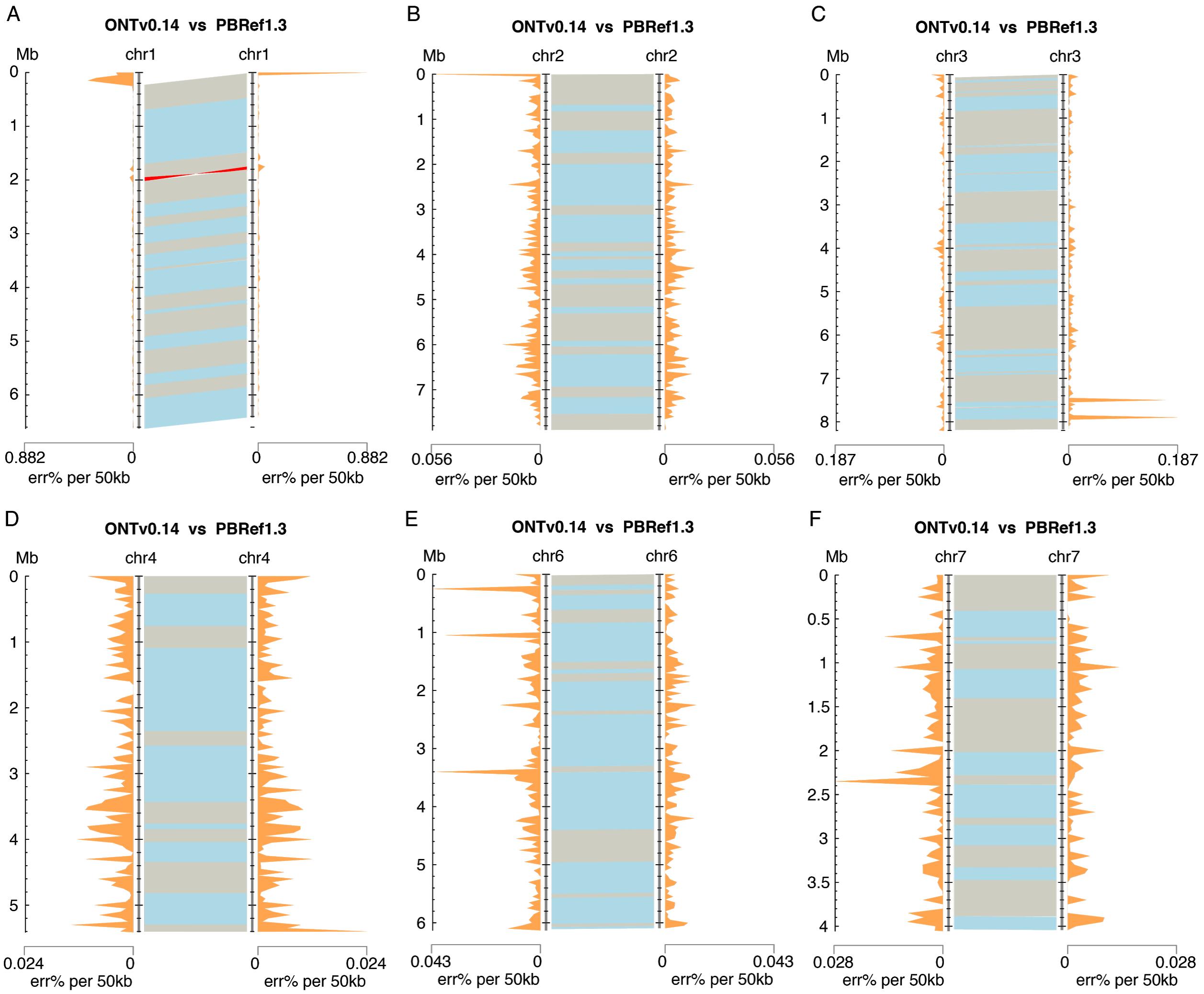


**Figure S6. Comparison of the two assemblies.** Comparison of each chromosome of 1, 2, 3, 4, 6, 7 between the genome assembly with Nanopore long reads (ONTv0.14) and the assembly with PacBio long reads (PBRef1.3). Distributions of percentages of errors (err%) per 50 kb are plotted separately along a chromosome of each assembly (orange shades). In between two assemblies, alignments using Nucmer are displayed, neighboring alignments are indicated by two colors. An alignment of chromosome 1 highlighted in red represents the sequence inverted in the two assemblies.

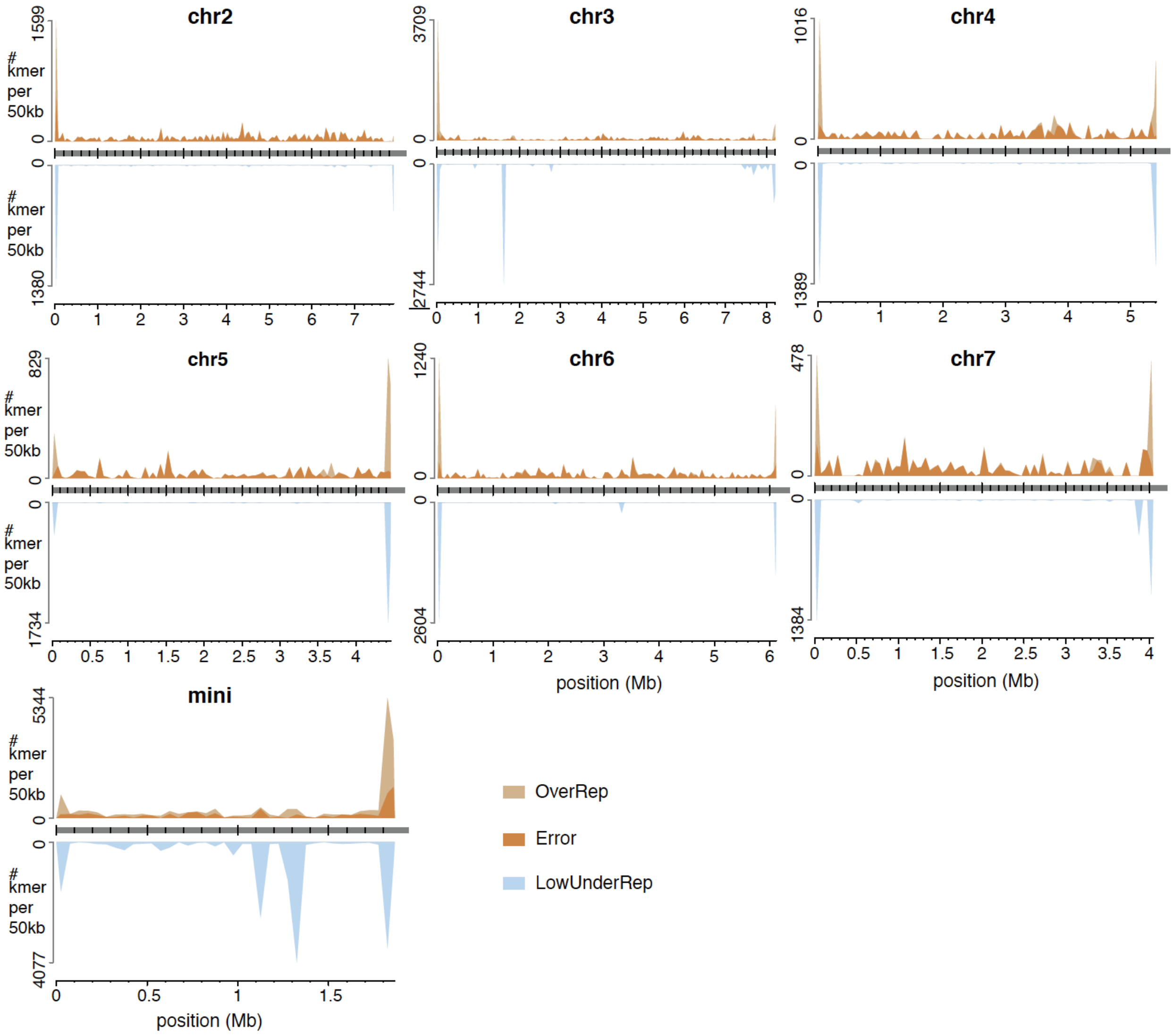


**Figure S7. KAD profiling identified errors in the fungal genome assembly.** Error k-mers and under-represented k-mers were mapped to the B71Ref1.5 assembly. Each k-mer was allowed to map to up to 100 locations. The number of k-mers in each group per 50 kb was determined and plotted versus the position of the 50-kb window in the assembly.

### 2. SUPPLEMENTARY TABLES

### Table S1. KAD evaluation for genomes with different sizes

| Species | Error rate | Total | Good | Error | OverRep | LowUnderRep | HighUnderRep |
|---|---|---|---|---|---|---|---|
| | 0.0% | 4548910 | 4539657 | 1800 | - | - | - |
| | 0.1% | 4663639 | 4427290 | 116529 | - | 107061 | - |
| | 0.2% | 4775534 | 4318087 | 228424 | - | 211601 | - |
| | 0.3% | 4884970 | 4210365 | 337808 | - | 314730 | - |
| | 0.4% | 4990815 | 4106588 | 443625 | - | 412599 | - |
| *Escherichia coli* (4.7M) | 0.5% | 5094366 | 4004985 | 547215 | - | 509625 | 4 |
| | 0.6% | 5196705 | 3904735 | 649523 | - | 605135 | 13 |
| | 0.7% | 5295171 | 3807855 | 747902 | - | 697906 | 8 |
| | 0.8% | 5393154 | 3711603 | 845982 | 1 | 790683 | 7 |
| | 0.9% | 5486257 | 3620195 | 938987 | - | 876633 | 67 |
| | 1.0% | 5581418 | 3527208 | 1034172 | - | 965057 | 65 |
| | 0.0% | 11534681 | 11532276 | 666 | 9 | - | - |
| | 0.1% | 11834109 | 11246470 | 300084 | 3 | 276458 | - |
| | 0.2% | 12126866 | 10967407 | 592677 | 4 | 547525 | - |
| | 0.3% | 12413823 | 10693829 | 879561 | 3 | 811664 | - |
| | 0.4% | 12691950 | 10430068 | 1157394 | 6 | 1067603 | 23 |
| *Saccharomyces cerevisiae* (12.4M) | 0.5% | 12962959 | 10171315 | 1428385 | 6 | 1320702 | 19 |
| | 0.6% | 13231843 | 9913145 | 1696896 | 35 | 1570246 | 41 |
| | 0.7% | 13490470 | 9667938 | 1954910 | 56 | 1807557 | 74 |
| | 0.8% | 13742581 | 9425712 | 2206906 | 48 | 2043548 | 115 |
| | 0.9% | 13989903 | 9190172 | 2453569 | 98 | 2270777 | 199 |
| | 1.0% | 14231805 | 8958320 | 2695625 | 51 | 2497430 | 246 |
| | 0.0% | 110396645 | 110383665 | 7581 | - | - | - |
| | 0.1% | 113333351 | 107628128 | 2943828 | 2 | 2607666 | 11 |
| | 0.2% | 116197118 | 104939324 | 5806447 | 26 | 5156811 | 50 |
| | 0.3% | 118995639 | 102312132 | 8602890 | 57 | 7648278 | 110 |
| | 0.4% | 121724901 | 99733757 | 11329648 | 195 | 10093016 | 309 |
| *Arabidopsis thaliana* (121.6M) | 0.5% | 124396465 | 97223330 | 13998745 | 294 | 12489859 | 510 |
| | 0.6% | 126992337 | 94775827 | 16590908 | 457 | 14816428 | 934 |
| | 0.7% | 129529463 | 92385334 | 19124826 | 586 | 17097108 | 1377 |
| | 0.8% | 132008176 | 90048493 | 21600197 | 818 | 19329670 | 2114 |
| | 0.9% | 134429001 | 87761673 | 24017435 | 889 | 21517555 | 2555 |
| | 1.0% | 136798533 | 85537682 | 26383078 | 1230 | 23644835 | 3615 |

| Species | Error rate | Total | Good | Error | OverRep | LowUnderRep | HighUnderRep |
|---|---|---|---|---|---|---|---|
| *Oryza sativa japonica* (381.3M) | 0.0% | 284649541 | 283973573 | 440792 | 376 | - | - |
| | 0.1% | 293048984 | 276986445 | 8837368 | 663 | 6469411 | 24 |
| | 0.2% | 301630653 | 269975265 | 17372275 | 4128 | 12797215 | 277 |
| | 0.3% | 310000648 | 263119089 | 25675007 | 11491 | 19001831 | 645 |
| | 0.4% | 318151197 | 256439194 | 33745604 | 21517 | 25082063 | 1579 |
| | 0.5% | 326115127 | 249894205 | 41616744 | 36063 | 31053713 | 2661 |
| | 0.6% | 333874466 | 243514123 | 49276240 | 51311 | 36898630 | 5125 |
| | 0.7% | 341429487 | 237318726 | 56728638 | 71246 | 42575832 | 7670 |
| | 0.8% | 348827968 | 231229232 | 64025654 | 91841 | 48189881 | 11027 |
| | 0.9% | 356027515 | 225274121 | 71119066 | 113871 | 53674385 | 16382 |
| | 1.0% | 363085818 | 219469933 | 78072025 | 137824 | 59028815 | 20380 |
| *Zea mays* (2.2G) | 0.0% | 694805059 | 694653026 | 96287 | 5 | - | - |
| | 0.1% | 731967220 | 674038296 | 36595238 | 94812 | 14254619 | 168 |
| | 0.2% | 767452397 | 654358872 | 70706856 | 424690 | 28163692 | 1571 |
| | 0.3% | 801657764 | 635575442 | 103184653 | 953034 | 41788648 | 4628 |
| | 0.4% | 834831023 | 617551401 | 134462119 | 1599602 | 55172779 | 11036 |
| | 0.5% | 867174926 | 600173658 | 164807838 | 2338867 | 68345201 | 20352 |
| | 0.6% | 898632235 | 583491214 | 194239297 | 3146387 | 81178580 | 33482 |
| | 0.7% | 929497520 | 567239832 | 223057749 | 3981201 | 93858306 | 51639 |
| | 0.8% | 959639627 | 551470062 | 251177075 | 4833109 | 106306417 | 75688 |
| | 0.9% | 989327920 | 536187985 | 278826613 | 5696910 | 118527640 | 104957 |
| | 1.0% | 1018359462 | 521262157 | 305899085 | 6552890 | 130555357 | 140866 |

**Table S2. Ratios of error k-mers detected by KAD to total simulated errors**

| Simulated Error Rate | Error k-mers/total simulated errors | | | | |
|---|---|---|---|---|---|
| | *E. coli* | Yeast | Arabidopsis | Rice | Maize |
| 0.1% | 25.1 | 24.7 | 24.6 | 23.6 | 17.1 |
| 0.2% | 24.6 | 24.4 | 24.3 | 23.2 | 16.6 |
| 0.3% | 24.3 | 24.1 | 24.0 | 22.9 | 16.1 |
| 0.4% | 23.9 | 23.8 | 23.7 | 22.6 | 15.7 |
| 0.5% | 23.6 | 23.5 | 23.4 | 22.3 | 15.4 |
| 0.6% | 23.3 | 23.3 | 23.1 | 22.0 | 15.2 |
| 0.7% | 23.0 | 23.0 | 22.8 | 21.7 | 14.9 |
| 0.8% | 22.8 | 22.7 | 22.6 | 21.4 | 14.7 |
| 0.9% | 22.5 | 22.4 | 22.3 | 21.1 | 14.5 |
| 1.0% | 22.3 | 22.2 | 22.0 | 20.9 | 14.3 |

**Table S3. TCRs and FCRs with different k-mer lengths in five genomes**

| k-mer type | k-mer size = 25bp | | k-mer size = 31bp | |
|---|---|---|---|---|
| | TCR | FCR | TCR | FCR |
| *E. coli* | 99.95% | 0.14% | 99.99% | 0.01% |
| Yeast | 99.77% | 0.02% | 99.98% | 0.03% |
| Arabidopsis | 99.29% | 0.02% | 99.98% | 0.02% |
| Rice | 94.73% | 0.03% | 97.20% | 0.05% |
| Maize | 67.73% | 0.43% | 79.98% | 0.30% |

Note: The genomes with 1% error rate and reads with 50x sequencing depth were simulated for KAD analyses.

**Table S4. KAD results for different k-mer lengths in maize**

| k-mer length | Total k-mer | Error k-mer | TCR | FCR |
|---|---|---|---|---|
| 19bp | 771458124 | 200423675 | 61.96% | 0.69% |
| 25bp | 1018359462 | 305899085 | 67.73% | 0.43% |
| 31bp | 1242304788 | 409947728 | 79.98% | 0.30% |
| 37bp | 1452463572 | 512773313 | 85.25% | 0.22% |
| 43bp | 1648425251 | 612332775 | 89.11% | 0.16% |
| 49bp | 1830091570 | 707462763 | 91.91% | 0.12% |
| 55bp | 1648425251 | 612332775 | 90.05% | 0.14% |

Note: The maize genome with 1% error rate and reads with 50x sequencing depth were simulated for KAD analyses.

### Table S5. KAD results for different types of simulated errors in the *E. coli* genome

| Variation Type | Variant Rate or Number | Total | Good | Error | OverRep | LowUnderRep | HighUnderRep |
|---|---|---|---|---|---|---|---|
| Single nucleotide substitution | 0% | 4,548,910 | 4,539,657 | 1,800 | 0 | - | - |
| | 1% | 5,578,487 | 3,529,182 | 1,031,131 | 0 | 963,312 | 128 |
| | 2% | 6,386,764 | 2,737,137 | 1,839,231 | 0 | 1,721,377 | 400 |
| | 3% | 7,020,149 | 2,115,799 | 2,472,653 | 2 | 2,318,318 | 1,040 |
| | 4% | 7,517,633 | 1,630,011 | 2,969,918 | 6 | 2,783,142 | 2,582 |
| | 5% | 7,899,276 | 1,257,537 | 3,351,604 | 20 | 3,140,719 | 4,357 |
| | 6% | 8,196,595 | 968,859 | 3,649,029 | 12 | 3,417,787 | 6,828 |
| | 7% | 8,436,939 | 734,217 | 3,889,440 | 4 | 3,641,208 | 8,602 |
| | 8% | 8,610,273 | 565,651 | 4,062,875 | 0 | 3,801,457 | 10,139 |
| | 9% | 8,748,265 | 431,448 | 4,200,955 | 0 | 3,930,649 | 11,402 |
| | 10% | 8,854,592 | 327,673 | 4,307,309 | 1 | 4,028,878 | 12,178 |
| Short Indel | 0% | 4,548,910 | 4,548,676 | 161 | 0 | - | - |
| | 1% | 5,564,369 | 3,552,865 | 1,015,401 | 0 | 981,033 | 63 |
| | 2% | 6,368,450 | 2,759,862 | 1,819,452 | 0 | 1,763,658 | 381 |
| | 3% | 6,998,264 | 2,142,379 | 2,449,010 | 26 | 2,375,067 | 1,255 |
| | 4% | 7,489,810 | 1,659,910 | 2,940,688 | 1 | 2,854,436 | 2,333 |
| | 5% | 7,881,345 | 1,276,817 | 3,332,252 | 0 | 3,233,063 | 4,259 |
| | 6% | 8,179,652 | 983,968 | 3,630,484 | 11 | 3,522,968 | 6,186 |
| | 7% | 8,416,771 | 757,355 | 3,867,753 | 1 | 3,746,155 | 7,646 |
| | 8% | 8,603,010 | 573,458 | 4,054,043 | 10 | 3,927,478 | 9,257 |
| | 9% | 8,740,454 | 435,846 | 4,191,587 | 4 | 4,062,813 | 10,874 |
| | 10% | 8,847,122 | 335,914 | 4,298,202 | 15 | 4,161,328 | 11,880 |
| Redundancy | 0 | 4,548,910 | 4,548,676 | 161 | 0 | - | - |
| | 50 | 5,564,369 | 3,552,865 | 1,015,401 | 0 | 981,033 | 63 |
| | 100 | 6,368,450 | 2,759,862 | 1,819,452 | 0 | 1,763,658 | 381 |
| | 150 | 6,998,264 | 2,142,379 | 2,449,010 | 26 | 2,375,067 | 1,255 |
| | 200 | 7,489,810 | 1,659,910 | 2,940,688 | 1 | 2,854,436 | 2,333 |
| | 250 | 7,881,345 | 1,276,817 | 3,332,252 | 0 | 3,233,063 | 4,259 |
| | 300 | 8,179,652 | 983,968 | 3,630,484 | 11 | 3,522,968 | 6,186 |
| | 350 | 8,416,771 | 757,355 | 3,867,753 | 1 | 3,746,155 | 7,646 |
| | 400 | 8,603,010 | 573,458 | 4,054,043 | 10 | 3,927,478 | 9,257 |
| | 450 | 8,740,454 | 435,846 | 4,191,587 | 4 | 4,062,813 | 10,874 |
| | 500 | 8,847,122 | 335,914 | 4,298,202 | 15 | 4,161,328 | 11,880 |
| Assembly Collapse | 0 | 4,548,910 | 4,539,657 | 1,800 | 0 | - | - |
| | 50 | 4,549,963 | 4,301,432 | 2,853 | 0 | 228,586 | 43 |
| | 100 | 4,551,074 | 4,002,122 | 3,964 | 0 | 512,834 | 5 |
| | 150 | 4,552,208 | 3,764,014 | 5,098 | 0 | 742,915 | 112 |
| | 200 | 4,553,141 | 3,541,773 | 6,031 | 0 | 953,956 | 30 |
| | 250 | 4,554,233 | 3,334,306 | 7,123 | 0 | 1,149,967 | 182 |
| | 300 | 4,555,473 | 3,048,873 | 8,363 | 0 | 1,420,813 | 327 |
| | 350 | 4,556,419 | 2,804,924 | 9,309 | 0 | 1,658,986 | 1,323 |
| | 400 | 4,557,569 | 2,585,461 | 10,459 | 0 | 1,868,337 | 431 |
| | 450 | 4,558,583 | 2,350,819 | 11,473 | 0 | 2,092,722 | 1,420 |
| | 500 | 4,559,592 | 2,068,398 | 12,482 | 0 | 2,362,767 | 1,444 |

**Table S6. Source of sequences of B71Ref1.5**

| seq | order | Source | Source start | Source end | strand | note |
|---|---|---|---|---|---|---|
| chr1 | 1 | ONTv0.14_ chr1 | 1 | 6627575 | plus | Nucmer between ONTv0.14&PBRef1.3 |
| chr2 | 1 | ONTv0.14_ chr2 | 1 | 8873 | plus | Nucmer between ONTv0.14&PBRef1.3 |
| chr2 | 2 | PBRef1.3_ chr2 | 1 | 7899709 | plus | Nucmer between ONTv0.14&PBRef1.3 |
| chr3 | 1 | ONTv0.14_ chr3 | 1 | 8206634 | plus | Nucmer between ONTv0.14&PBRef1.3 |
| chr4 | 1 | ONTv0.14_ chr4 | 1 | 5412286 | plus | Nucmer between ONTv0.14&PBRef1.3 |
| chr5 | 1 | ONTv0.14_ chr5 | 1 | 4459566 | plus | Nucmer between ONTv0.14&PBRef1.3 |
| chr6 | 1 | ONTv0.14_ chr6 | 1 | 18307 | plus | Nucmer between ONTv0.14&PBRef1.3 |
| chr6 | 2 | PBRef1.3_ chr6 | 1 | 6092616 | plus | Nucmer between ONTv0.14&PBRef1.3 |
| chr6 | 3 | ONTv0.14_ chr6 | 6123809 | 6131042 | plus | Nucmer between ONTv0.14&PBRef1.3 |
| chr7 | 1 | ONTv0.14_ chr7 | 1 | 158 | plus | Nucmer between ONTv0.14&PBRef1.3 |
| chr7 | 2 | PBRef1.3_ chr7 | 1 | 4043141 | plus | Nucmer between ONTv0.14&PBRef1.3 |
| chr7 | 3 | ONTv0.14_ chr7 | 4052747 | 4056003 | plus | Nucmer between ONTv0.14&PBRef1.3 |
| mini | 1 | ONTv0.14_ mini | 2921 | 1868262 | plus | Nucmer between ONTv0.14&PBRef1.3 (2,920 bp were trimmed from mini due to the extra sequence beyond telomere sequences) |
| mt | 1 | PBRef1.3_ mt | 1 | 34996 | plus | Nucmer between ONTv0.14&PBRef1.3 |

**Table S7. KAD summary of B71Ref1.5**

| Assembly | B71Ref1.5 |
|---|---|
| Total | 45,176,684 |
| Good | 44,390,516 |
| Error | 51,367* |
| OverRep | 4,983 |
| LowUnderRep | 32,437 |
| HighUnderRep | 42,485 |

* indicated ~2,568 (51,367/20) base errors that used 20 as the conversion rate.
